# Convis: A Toolbox To Fit and Simulate Filter-based Models of Early Visual Processing

**DOI:** 10.1101/169284

**Authors:** Jacob Huth, Timothée Masquelier, Angelo Arleo

**Keywords:** vision model toolbox, retina model, primary visual cortex model, Python, GPU, Theano, PyTorch

## Abstract

We developed *Convis*, a Python simulation toolbox for large scale neural populations which offers arbitrary receptive fields by 3D convolutions executed on a graphics card. The resulting software proves to be flexible and easily extensible in Python, while building on the PyTorch library [32], which was previously used successfully in deep learning applications, for just-in-time optimization and compilation of the model onto CPU or GPU architectures. An alternative implementation based on Theano [33] is also available, although not fully supported.

Through automatic differentiation, any parameter of a specified model can be optimized to approach a desired output which is a significant improvement over e.g. Monte Carlo or particle optimizations without gradients. We show that a number of models including even complex non-linearities such as contrast gain control and spiking mechanisms can be implemented easily.

We show in this paper that we can in particular recreate the simulation results of a popular retina simulation software VirtualRetina [35], with the added benefit of providing (1) arbitrary linear filters instead of the product of Gaussian and exponential filters and (2) optimization routines utilizing the gradients of the model. We demonstrate the utility of 3d convolution filters with a simple direction selective filter. Also we show that it is possible to optimize the input for a certain goal, rather than the parameters, which can aid the design of experiments as well as closed-loop online stimulus generation. Yet, Convis is more than a retina simulator. For instance it can also predict the response of V1 orientation selective cells.

Convis is open source under the GPL-3.0 license and available from https://github.com/jahuth/convis/ with documentation at https://jahuth.github.io/convis/.

## 2 Introduction

We developed Convis as an extension to the popular PyTorch toolbox [32] that can implement models of responses to visual inputs on a large scale and, using automatic differentiation, derive the gradient of the output with respect to any input parameter. Linear filtering is done as convolutions, giving full flexibility in the shape and temporal structure of receptive fields. As an example, we show the utility of this method by examining models of retinal ganglion cells (RGCs), ranging from a very simple linear-non-linear model to a many-parameter mechanistic model which includes recurrent contrast gain control. Further, we show that this toolbox can simulate cell populations on a similar scale and in comparable time to previous large-scale retina simulators, even though the computations refrain from simplifying the shape of receptive fields.

We aim to bridge two use cases of vision models which tended use distinct software solutions in the past. To understand the structures and processes of the visual system, eg. how plasticity shapes representations[22], theorists are interested in a relatively large portion of the visual field, which requires a large number of neurons to be simulated according to our current understanding of the typical response of the respective cells. Experimentalists are in contrast concerned with the responses of a set of particular, very individual cells. While the mechanisms of the model used are often similar, the theorist requires a large number of cells to be simulated rapidly and efficiently with identical parameters from the literature and the experimentalist needs a method to adjust model parameters such that the model recreates the behavior of the cells observed in experiments.

The toolbox we created can be used in these two applications.

### 2.1 Population Simulation Software

#### 2.1.1 Large Scale Retina Models

The retina is the first stage of neural computation in vision. Models of different complexity exist that predict retinal ganglion cell responses for a wide range of ganglion cell types (from simple ON and OFF alpha cells up to direction selective cells). Some efforts were made to create large scale population simulators (eg. VirtualRetina [35] and COREM [20]) that can create the responses of thousands of retinal ganglion cells with efficient computations. These can be used as input to higher visual functions which generally need a large number of cells as input.

However, these simulators are not very useful for experimentalists who want to model the behavior of cells recorded in their experiments. The parameters of the model cannot be directly inferred from an experiment and fitting the simulation to the data can only be done via parameter grid search or Monte Carlo methods - both being very computation intense. Retinal processing is - in contrast to the rest of the visual system - easily accessible to multi-array recordings. Despite the relative consistency of retinal circuitry, a high number of ganglion-cell response classes exists [21]. To characterize the properties of each class of cells, linear-non-linear cascade models are the current gold standard. However, the properties of eg. contrast gain control are not easily extracted from data: Experiments usually explore only distinct levels of contrast, for instance [12]. [29] proposed a mechanism for contrast gain control, which accounts for the phase advance and transfer function recorded in RGC cells. It is incorporated into large scale retina population simulators such as VirtualRetina in a formulation that takes local contrast, rather than global contrast, into account.

To keep computations tractable, VirtualRetina uses recursive filtering rather than actual convolution, which limits the possible applications of these models to circular receptive fields and exponential temporal filters. But receptive fields are rarely circular, in most studies RGC receptive fields will be visualized as tilted ellipses (eg. [16]), while [13] even show that the irregular shape of receptive fields is linked to their objective to tile the visual field. To incorporate non-circular receptive fields, Convis allows for recursive as well as dense numerical filters, which are convolved with the input on a graphics card. Cell responses with detailed receptive fields can thus be simulated efficiently and the model can be fitted to experimental recordings since the gradients of the computations are available.

#### 2.1.2 Computations in the Retina

RGCs transmit the output of the retina to different brain regions, mainly the LGN. It is estimated that there exist at least 30 different RGC types [28], each population forming a tiling coverage of the visual space, responding uniquely to light stimuli and possesses unique proteins making it possible to distinguish between different types histochemically. The visual system, even at this low level, is complex. A model that predicts, for a given set of stimuli, the corresponding responses accurately might not be able to do so for some novel stimuli, due to over-fitting. The responses to “natural images” lead to more complex models than the responses to high-luminance on-off pulses, moving bars and gratings or even random checkerboard stimuli [31]. Still, a “natural stimulus” will also not cover all modes of behavior and even more complex models are conceivable. Choosing the complexity of a model is a hard problem in itself. The model should explain the data while being as simple as possible to avoid over-fitting as data is always limited. The simplest approximation of RGC responses can be obtained with a linear-non-linear model that predicts the firing probability of a cell by multiplying the input with a spatial or spatio-temporal receptive field and then applying a non-linearity [18, 15]. When a stimulus is spatially more complex and has a finer resolution, it becomes apparent that subunits exist that integrate the stimulus independently under a first non-linearity before being summed and subject to a second non-linearity. Responses to these stimuli can be predicted with subunit models (also called LN-cascade models as they stack multiple levels of linear and non-linear processing). We implemented a range of models on this spectrum of complexity and assessed whether parameters could be estimated efficiently.

#### 2.1.3 A Complex Retina Model: VirtualRetina

As an example of what Convis can do, we implemented a retina model close to the mathematical definition of VirtualRetina [35], but replaced the recursive method to compute linear filters with convolution operations.

The recursive definition of filters makes VirtualRetina very fast and efficient. Two-dimensional Gaussians for instance are computed independently for the x and y direction, such that the time complexity is scaling essentially linear with each dimension. But recursive filters have the drawback that their shape is very limited: the x-y separability is a huge constraint for receptive fields. A radially symmetric filter is a good approximation for a diffusion process, but neural connections tend to be sparser and more selective [13]. To simulate arbitrary, non-round, non-Gaussian and non-continuous spatio-temporal filters we used 1d, 2d or 3d convolutions, which have the drawback of being very inefficient when implemented in single threaded programs. To counter this flaw and to keep the code comprehensible and maintainable, we used the PyTorch library to create models and run optimized code on a GPU. Spatio-temporal filters with finite impulse responses can be implemented effectively with either a recursive definition that keeps N previous states or higher moments in memory (as used in VirtualRetina and COREM) or as a convolution with a 3D kernel. We go further into the advantages and disadvantages of the two methods in section 3.3.3. Although a convolution is computationally costly because it has to calculate the crossproduct of filter and image at each position of the 3D image, the use of GPUs for cheap parallel computations can speed up this process dramatically.

Five main types of processing happen in the layers of the retina [35]. Through the absorption of light in photoreceptors, the input is low-pass filtered in the spatial and temporal domain and the sensitivity of each photoreceptor is adjusted by adaptation mechanisms, eg. photopigment bleaching. Through the inhibitory surround signal of horizontal cells, luminance information is mostly removed in favour of contrast information. The photoreceptors already adapt to luminance to some degree, but they are quite slow (between seconds and minutes). The inhibition from the surround is much faster path of luminance invariance. The combination of short term adaptation to luminance and the center-surround receptive field can be modeled as spatio-temporal high pass filters. The weight of center and surround gives a cell a more phasic or tonic response to luminance. When both receptive fields are balanced, luminance is removed perfectly. This contrast signal is again gain controlled by Bipolar-Amacrine synapses, resulting in characteristic phase advance of contrast gain control [30] and rectified due to strictly positive synapses. Lastly, spikes are generated in ganglion cells and sent along their axons in the optic nerve.

Each ganglion cell type differs in the parameters of these operations. Each filter operation can react differently to spatial and temporal frequencies. While the simplest possible model (a linear-non-linear model) can capture the general shape of the classic, excitatory receptive field under one specific condition, changes in illumination and surrounding contrast as well as the presentation of natural images require non-linear adaptation mechanisms for both[27]. [14] showed that generalized linear models with scalar non-linearities fail to capture RGC responses to natural images. [26] created a systematic hierarchy of RGC models, starting with a linear-nonlinear model and improving performance by adding subunits to the model, which compute independent non-linear functions before their output is aggregated and again subjected to a non-linearity. They further improved performance with feedback mechanisms.

Consequently, RGC responses are most accurately modelled as a cascade linear-non-linear model with gain control. The stages of the VirtualRetina simulator for example consist of a linear, spatio-temporally non-separable filtering mimicking the Outer Plexiform Layer (OPL) which converts luminance into a contrast image via a center-surround receptive field. The second stage, modeling the contrast gain control properties of bipolar cells subject to shunting inhibition, is a leaky integrator with leak conductance that depends on a spatio-temporal neighborhood of contrast magnitude. The last stage creates spikes by applying a static non-linearity to the output of the bipolar cells and then using a Leaky Integrate and Fire model to convert current into precise spike times. The contrast gain control as implemented in VirtualRetina has two effects in retinal processing: (1) it filters frequencies and (2) a phase advance is applied to the signal. This is typically assessed by exposing the system to a sum of sine waves and varying the contrast. Not only does the gain change, leading to sub-linear output, but with increasing contrast the phase advance decreases the response time compared with lower contrast stimuli.

Fitting the VirtualRetina model completely from experimental data is not feasible. Many of the parameters have correlated effects, the number of overall parameters is high and the nonlinear nature of some parameters increases the difficulty further. Thus, the parameters used for simulations are usually taken from the literature (such as the average receptive field size and size of the suppressive surround) and remaining parameters are tuned with Monte Carlo optimization to achieve the desired firing rates [35].

The reimplementation of VirtualRetina in the Convis toolbox can in contrast be fitted to data by using the gradients of the parameters. The first gradients give information about the location of a nearby minimum (see Section 4.3), while the second derivatives can give information about the shape of the error function, specifically if there is a global minimum, and the interaction of parameters by calculating a Hessian Matrix (see Section 4.4).

### 2.2 Fitting Experimental Data

Experimentalists analyze the behaviour of retinal ganglion cells eg. by calculating the spike-triggered average (STA) and spike-triggered covariances (STC), which can then be used as receptive fields in generative linear-nonlinear models [16]. If more complex parameters are added to the model, more elaborate parameter search has to be performed (eg. Polak-Ribière in the supplementary material of [26]).

More complex models have to be fitted with very elaborate optimization procedures, usually by evaluating the model using many different sets of parameters, eg. by grid search (which is not feasible for even moderate parameter spaces), Monte-Carlo methods [8] or even genetic algorithms[10].

Recently [3] used the approach of multi-layer recurrent neural networks to fit RGCs. While the model can predict the responses in the data well, the model is not interpretable and not mechanistic. So their final model is a GLM-RNN hybrid in which the spatial filter is linear and the temporal dynamics are achieved with two reservoirs of recurrently connected neurons to capture many of the non-linear effects. While their general approach is very different from ours, they also use a computational framework (in their case Theano) that provided them with gradients to guide their optimization routine.

All of the three use cases can benefit from the Convis framework: We offer LN-cascade models for experimentalists with many optimization algorithms out of the box. For more complex, mechanistic and bio-inspired models, Convis provides a way to use gradient-guided rather than brute force optimization techniques. And finally experimental combinations of modelling and machine learning can draw on the models provided by Convis and the wealth of available PyTorch packages to create new approaches rapidly with a small code base.

## 3 Methods

### 3.1 Usage

#### 3.1.1 General Usability

We recommend using jupyter notebooks or IPython console to work with Convis [25]. This way, models can be built interactively, auto completion is available and data can be inspected directly and plotted.

For code examples we provide, we assume that the necessary packages are installed and a python interpreter is running with the following code already executed:

~~~
import numpy as np
import matplotlib.pylab as plt
import torch
import convis
~~~

Convis aims to be compatible with Python 2.7 and Python 3.6. Installation instructions and a more in depth documentation of features and usage can be found at https://jahuth.github.io/convis/.

#### 3.1.2 Creating and Running a Model

Models can be created by instantiating an instance of one of the model classes available in Convis, eg. the `convis.retina.Retina()` model or the models found in `convis.models`.*. Keeping with PyTorch, each model can be called as a function to process some input. Additionally, a run method can be called with an argument dt to automatically break down the input into chunks of length dt and processing one after another to limit memory usage. This code sample will run the Retina model on a moving grating stimulus:

~~~
retina = convis.retina.Retina() # creating a model
some_input = convis.samples.moving_grating(t=2000,x=20,y=20)
o = retina(some_input[0:100,:,:])
# or
o = retina.run(some_input,dt=100)
~~~

> By default the model will generate on and off spiking responses.
>
> Creating a simple LN model can be defined like this:

~~~
m = convis.models.LN()
m.conv.set_weight(np.random.rand(10,5,5))
~~~

Models similar to the definitions from [26] are already implemented: LN, LNLN (called LNSN for subunit), LNLN with feedback at the second stage (LNSNF) and both stages (LNFSNF) and LNLN models with delayed feedback (LNFDSNF). In contrast to Real et al., where the stimulus had only one dimension in space, our filters are 3D spatio-temporal filters.

#### 3.1.3 Streams

In our use case we deal with three-dimensional movies or five-dimensional tensors (including two additional dimensions for batches and color channels). But the input might be larger than the available memory or even infinite in duration if we stream input for instance from a webcam. Thus, we only compute the response to a time slice at a time, the size of which depends on the available graphics card memory and image dimensions. Normally, this would mean that each time slice would be computed independently and the history of each process is lost. To enable dynamics that have a longer reaction time than one time slice, we use state variables, eg. the values of a differential equation at a certain point in time and a section of the input for convolution operations. The downside of this method is that the time slices can not be computed in parallel, since they depend on the history of previous slices. Another downsize is that the computational graph will either grow infinitely, or has to be cut off at some point between time slices. The default solution is to cut the graph regularly, which can limit the possibility of fitting processes with very slow time constants.

Convis supports multiple input and output formats, such as sequences of images, videos,.npy and.inr files and input via a network connection. Internally these are all represented by the same interface as stream objects from which a number of frames can be taken and to which a number of frames can be appended. They can be automatically resampled to different frame rates (to match the discrete time step of the model) and if more than one input is required multiple streams can be synchronized using timestamps.

To ease handling in- and outputs, image streams and a Runner object can be used to automatically feed input into models.

~~~
inp = convis.streams.RandomStream(size=(10,10),level=0.2,mean=0.5)
# keeping the output in memory:
out1 = convis.streams.SequenceStream(sequence=np.ones((0,10,10)))
# visualizing the output using Tkinter:
out2 = convis.streams.run_visualizer()
# saving the output uncompressed to disc:
out3 = convis.streams.InrImageStreamWriter(’output.inr’)
# saving the output compressed to disc:
out4 = convis.streams.HDF5Writer(‘output.h5’)
~~~

If OpenCV is installed, a number of video files can be read from and written to and webcams can be accessed.

A Runner object ties together a model, an input stream and an output stream. The Runner can spawn its own thread that will constantly consume data from the input stream (if available), process the data and feed the output to the output stream to eg. visualize the data or save it to a file.

~~~
runner = convis.Runner(retina, input = inp, output = out4)
runner.start()
# … some time later
runner.stop()
~~~

To enable color processing or other additional channel information, such as RGC types, we pass input from one layer to the next as 5d tensors rather than 3d. We follow a convention for 3D convolution, which is to add one dimension for “batches”, which are independent and can be processed in parallel, and one dimension for “channels”, such as color. We add those dimensions as dimension 0 and 1, consistent with the dimensions used by PyTorch 3d convolution. An input with a set number of color channels requires the convolution filter to have the exact same number of input channels and the output of all the individual channel filters will be summed. Conversely a single filter can also generate multiple output channels, which can be passed to the next layer.

As an example the following filter accepts input with 3 channels and produces output of 3 channels, switching the red, green and blue channels:

~~~
m = convis.models.LN()
kernel = np.zeros((3,3,1,1,1))
kernel[0,1,:,:,:] = 1.0
kernel[1,2,:,:,:] = 1.0
kernel[2,0,:,:,:] = 1.0
m.conv.set_weight(kernel)
~~~

#### 3.1.4 Automatic Optimization

Using an optimization algorithm to fit a PyTorch model to data can be done by creating the appropriate Optimizer object, filling the gradient buffers and executing an optimization step:

~~~
x,y = convis.samples.generate_sample_data()
# variables x and y are 3d time series
m = convis.models.LN()
opt = torch.optim.Adam(m.parameters())
opt.zero_grad()
model_output = m(x)
loss = ((model_output[0] - y)**2).mean()
loss.backward(retain_graph=True)
opt.step()
~~~

We added the set optimizer and optimize methods to all Convis models to simplify this code:

~~~
x,y = convis.samples.generate_sample_data()
m = convis.models.LN()
m.set_optimizer.Adam(m.parameters())
m.optimize(x,y) # fit the model, such that input x will result in output y
~~~

The list and properties of all optimizers can be found in the PyTorch documentation (http://pytorch.org/docs/master/optim.html) or by using Tab-completion on `m.set optimizer.<tab>` and the `help()` function (see also Section 3.2.1).

### 3.2 Theory

#### 3.2.1 The Computational Graph and Automatic Optimization

A computational graph can represent mathematical operations on data by combining symbols for variables and operations in a an acyclic graph. In deep learning frameworks, such as Theano, Tensorflow or PyTorch, this graph is used to automatically derive gradients by applying back-propagation rules.

In an earlier iteration of the toolbox, which is still available on github https://github.com/jahuth/convis_theano, we used *Theano* [33] to create models that are fully implemented as computational graphs. In the current version, which uses PyTorch, the computational graph is build during computation and only used for automatically deriving the gradients, while the forward computations are just-in-time compiled either to for GPU or CPU. When the backward pass is used on an output variable, the graph is traversed backwards and the backwards computation of each operation is applied and used for back-propagation. The resulting values get added to the grad buffer of each variable along the way until the input variables are reached.

Adapting the parameters of a model is a lot easier when gradients are available. A small example is provided in Figure 7 where the response of a target exponential filter to a random event train is approximated by a second exponential filter, and Figure 8 where a receptive field was recovered by approximating the response of the first filter.

**Figure 1:**
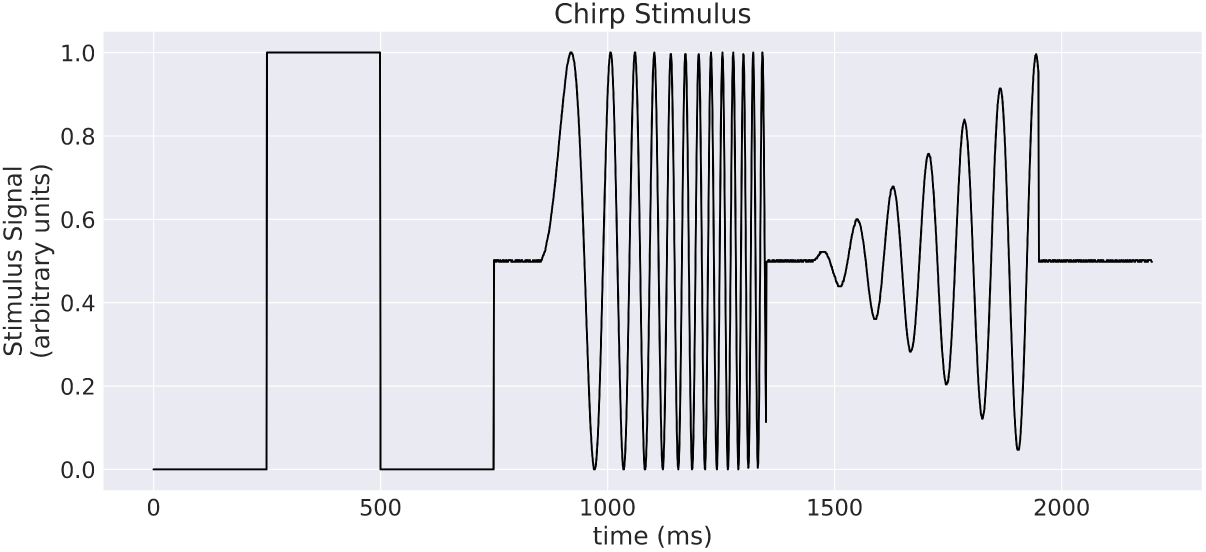
A “chirp” stimulus: the stimulus is comprised of an Off-On-Off pulse, then oscillations with increase in frequency, then oscillations with increase in amplitude.

**Figure 2:**
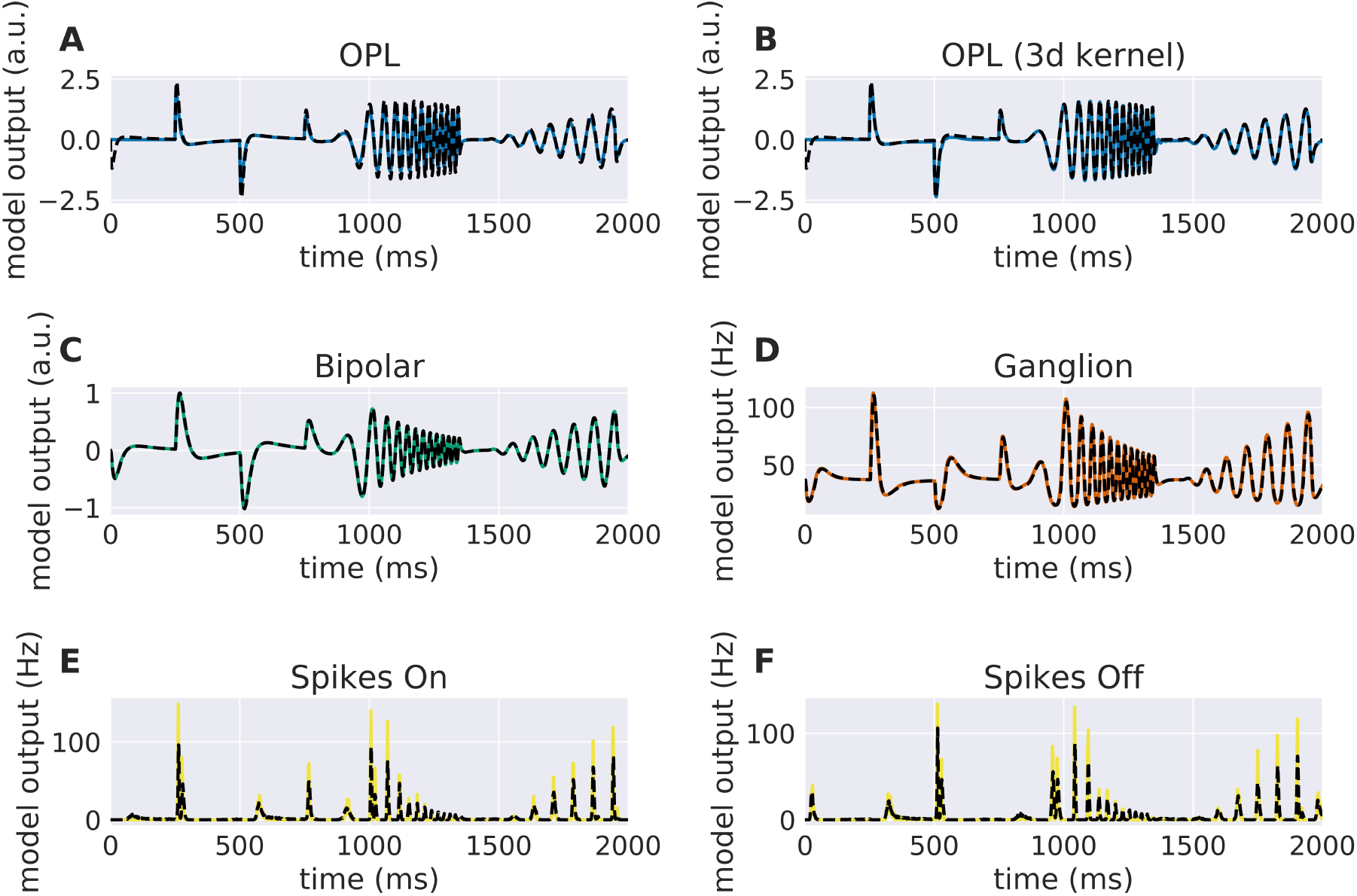
Comparison of Virtual Retina stages and the convolutional retina model to a chirp stimulus (see Figure 1). The response of Virtual Retina is shown as a dashed black line, the response of the Convis retina is shown as a solid lines of different colors. (A) shows the response of the linear OPL filter as configured with the same configuration as VirtualRetina, while (B) shows the response of a single 3d filter that was fitted to the desired output. (C-F) show the responses of the subsequent layers up to the spike generation. A quantification of the match between the lines can be seen in Figure 3. More detailed plots for each stage can be found in the additional material in Figures 11 to 15.

**Figure 3:**
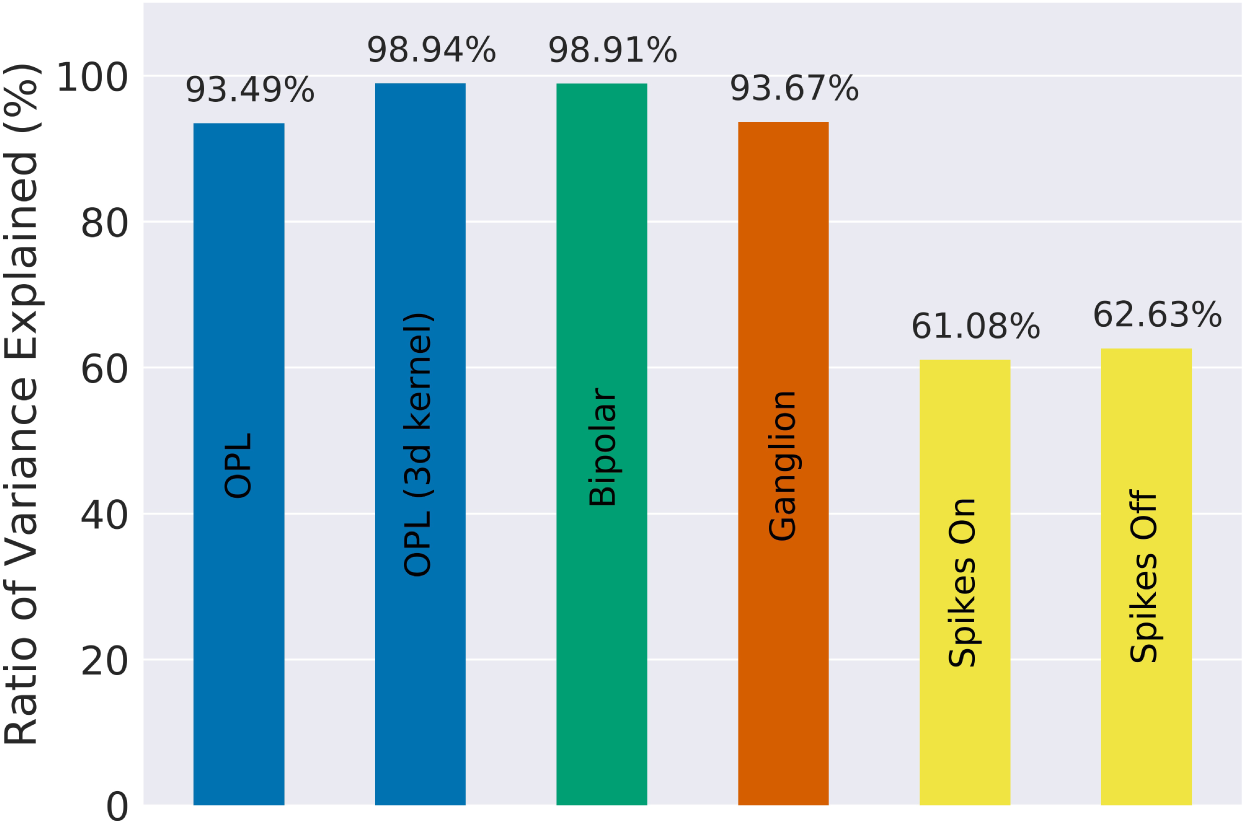
The fraction of variance of the original VirtualRetina explained for each individual stage of the model for the comparison simulations in Figure 2. Due to the filter resolution and numerical errors each stage has a slightly different trajectory than the original model. The spike generation explains a very small percentage of variance, as the process is inherently noisy. In Figure 2 For a more detailed comparison see Figures 11 to 15

**Figure 4:**
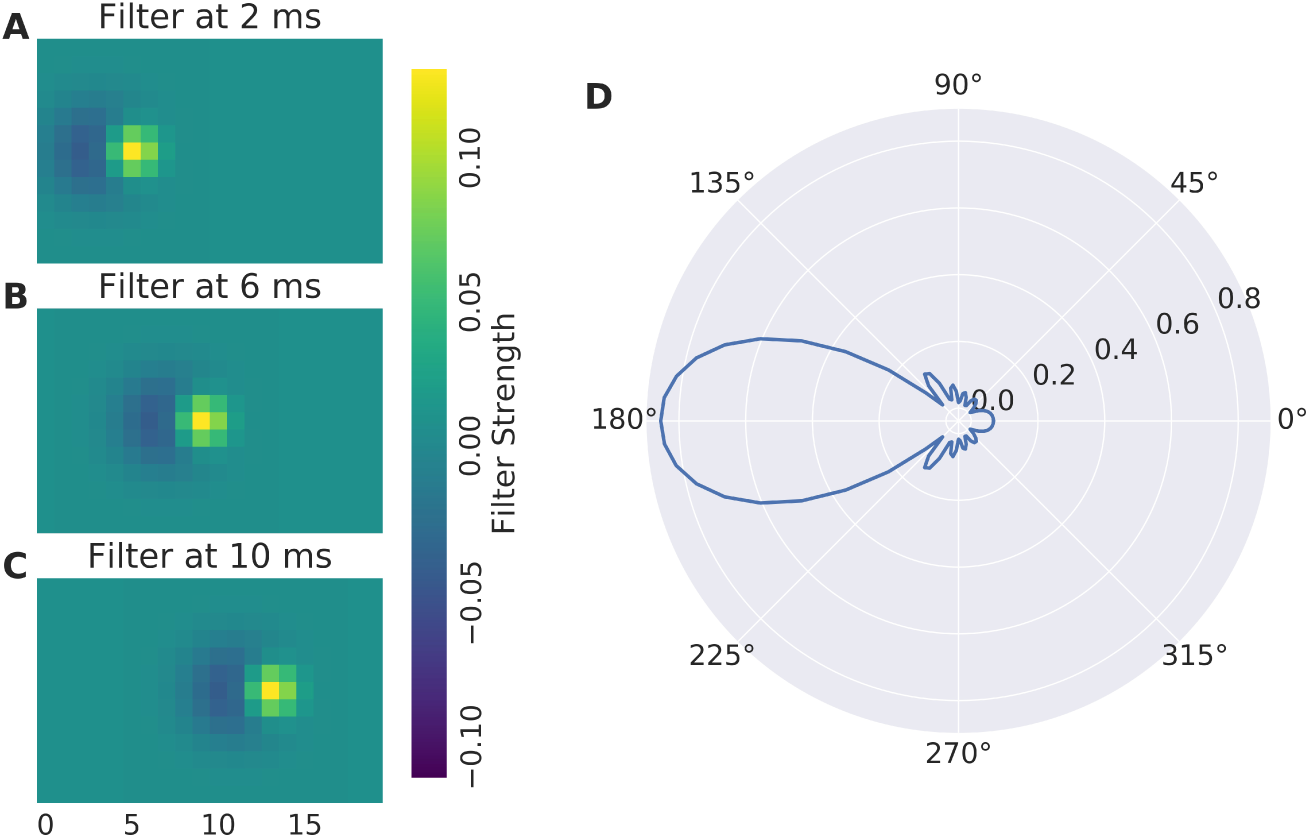
(A-C) Direction Selective Receptive Field and (D) direction tuning of a DS cell. Concentric rings are normalized response.

**Figure 5:**
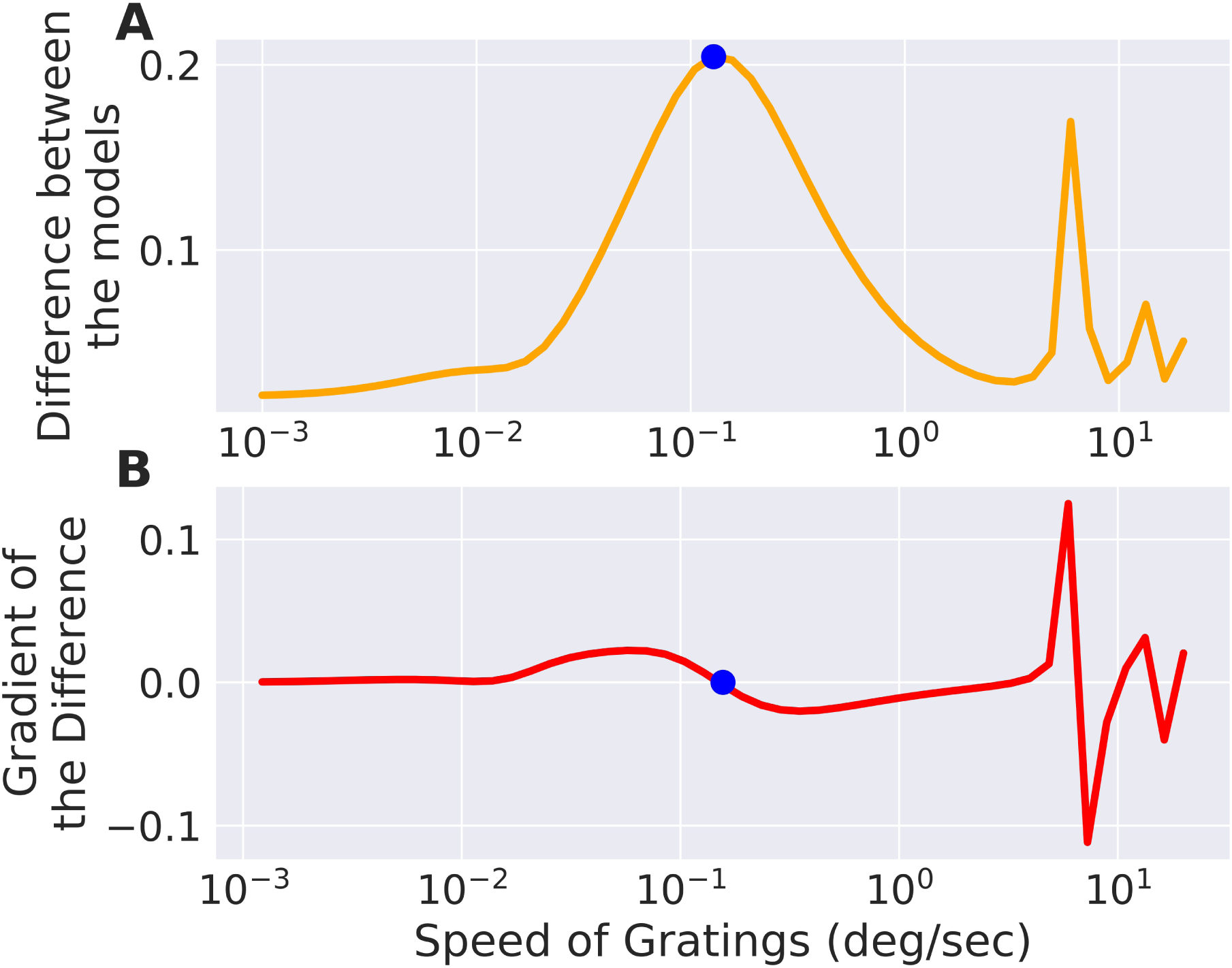
The difference between two models, slightly differing in their parameters, is low for very fast or very slow moving gratings, but in between there exists an optimum which is marked with a point. The same point is marked in the gradient of the error with respect to the grating speed parameter where it crosses 0. There are also secondary local maxima due to the periodicity of the stimulus.

**Figure 6:**
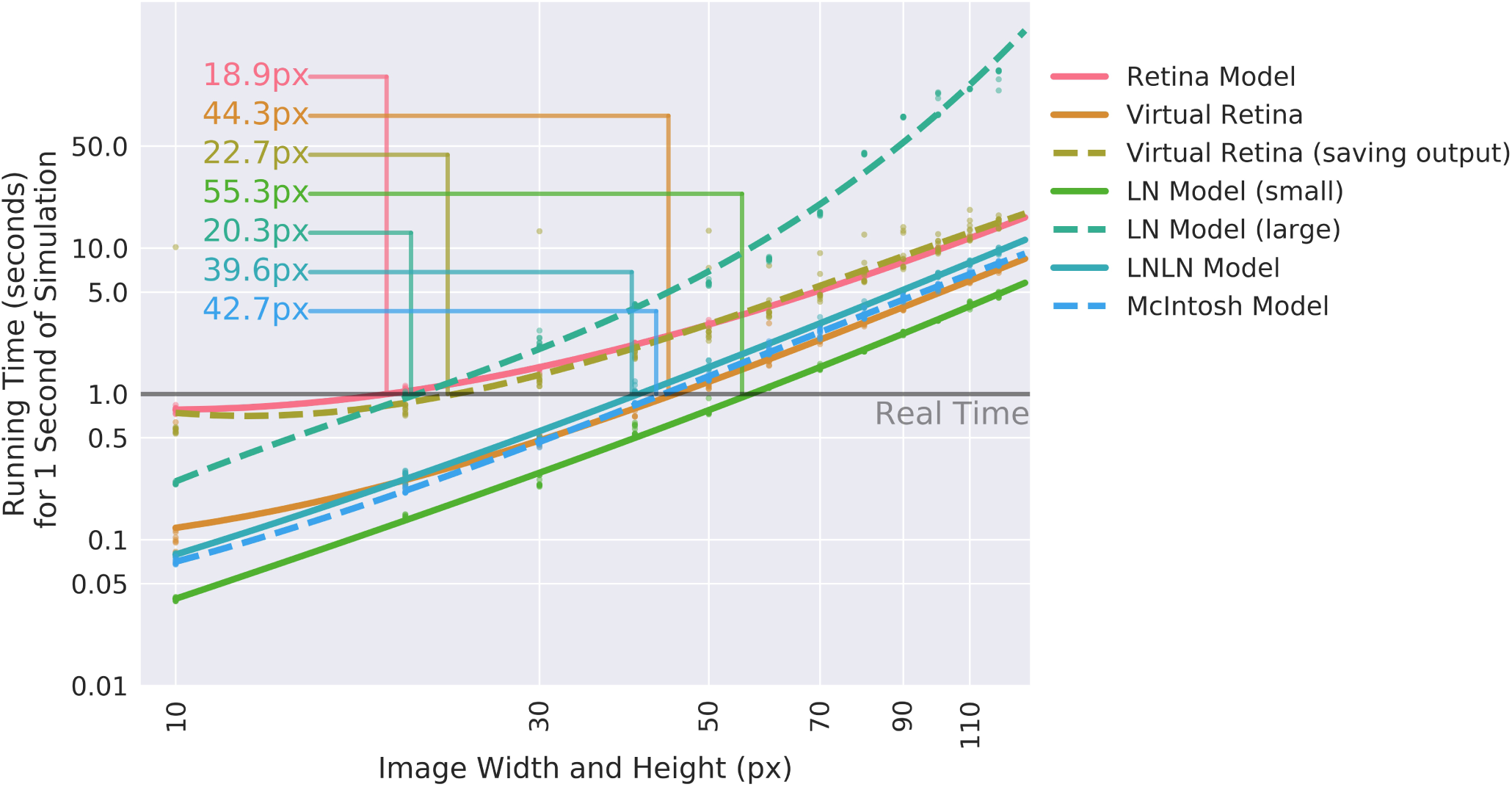
Comparison of the calculation speed of different Convis models and VirtualRetina [35] over a range of stimulus sizes. The stimuli were square videos of moving bars and the calculations were repeated 10 times. Dashed lines show when models cross from faster-than-real-time to slower-than-real-time and the approximate image size below which real-time processing is possible.

**Figure 7:**
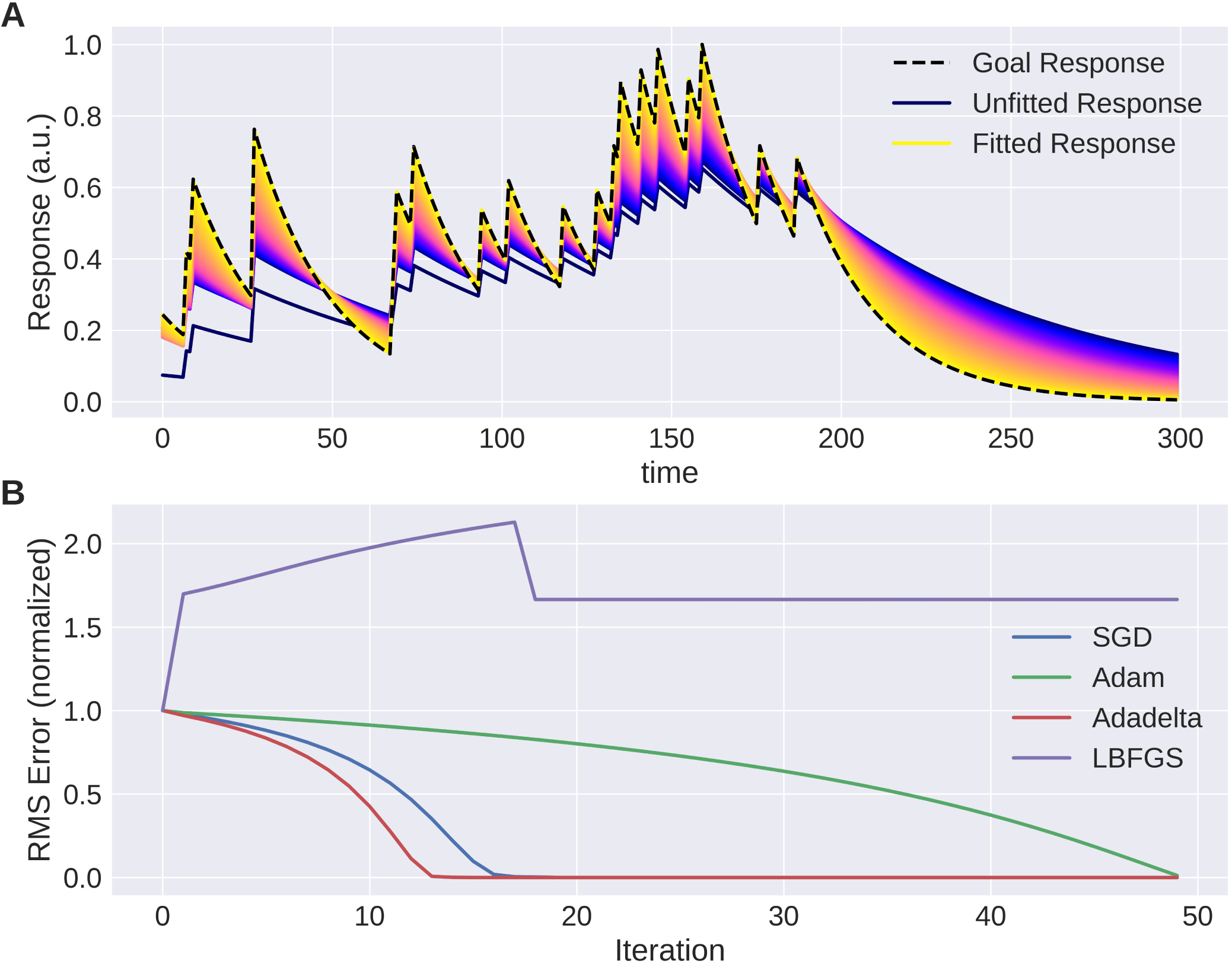
The trace of an exponential filter being fitted to a target: Top shows the different traces(with a very small adaptation between each iteration) from dark blue (beginning) to yellow (end). The bottom plot shows the instantaneous error over time for each trial. Note that using naive gradient descent the convergence slows down the closer we are to the solution.

**Figure 8:**
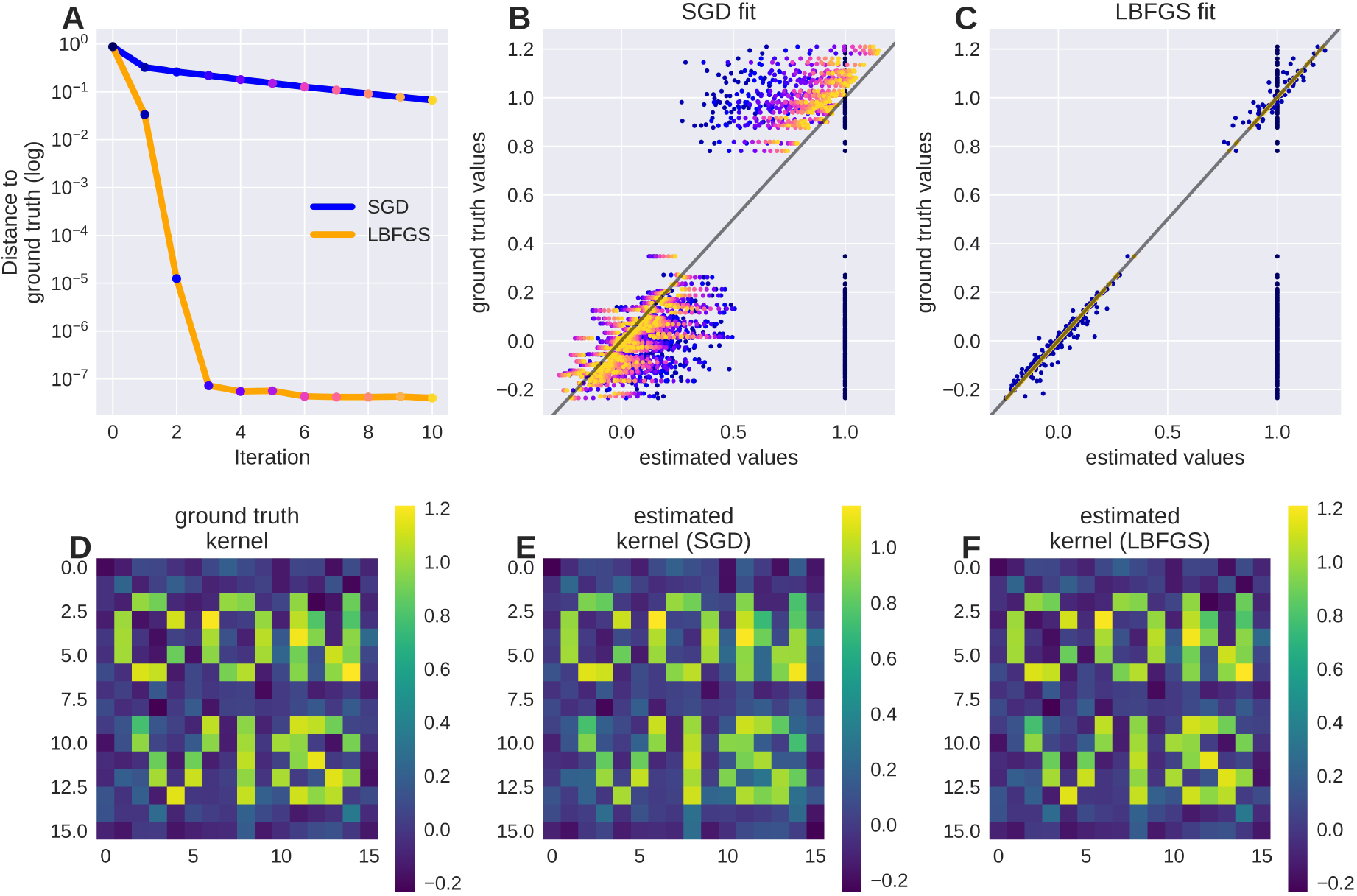
A receptive field of a simple linear-nonlinear model recovered using the gradients: Although the parameter space is large (16x16 pixel), the direction for each pixel can be estimated well. (A) shows the successive parameter values (from blue to yellow) compared to their ground truth when optimizing using Stochastic Gradient Descent, (B) shows the convergence of LBFGS optimization which finds the perfect solution in only one step; (C) shows the original filter, a sparse text sample with multiplicative noise and (D) shows the recovered filter.

PyTorch offers a range of optimization algorithms [32]. Some examples are: Sochastic Gradient Descent, Adam/Adamx (as used in [3]), Limited-memory BroydenFletcherGoldfarbShanno (LBFGS), RMSProp. The BroydenFletcherGoldfarbShanno algorithm is a quasi-Newton method and works very well for a large number of almost linear-behaving parameters (see Figure 9).

**Figure 9:**
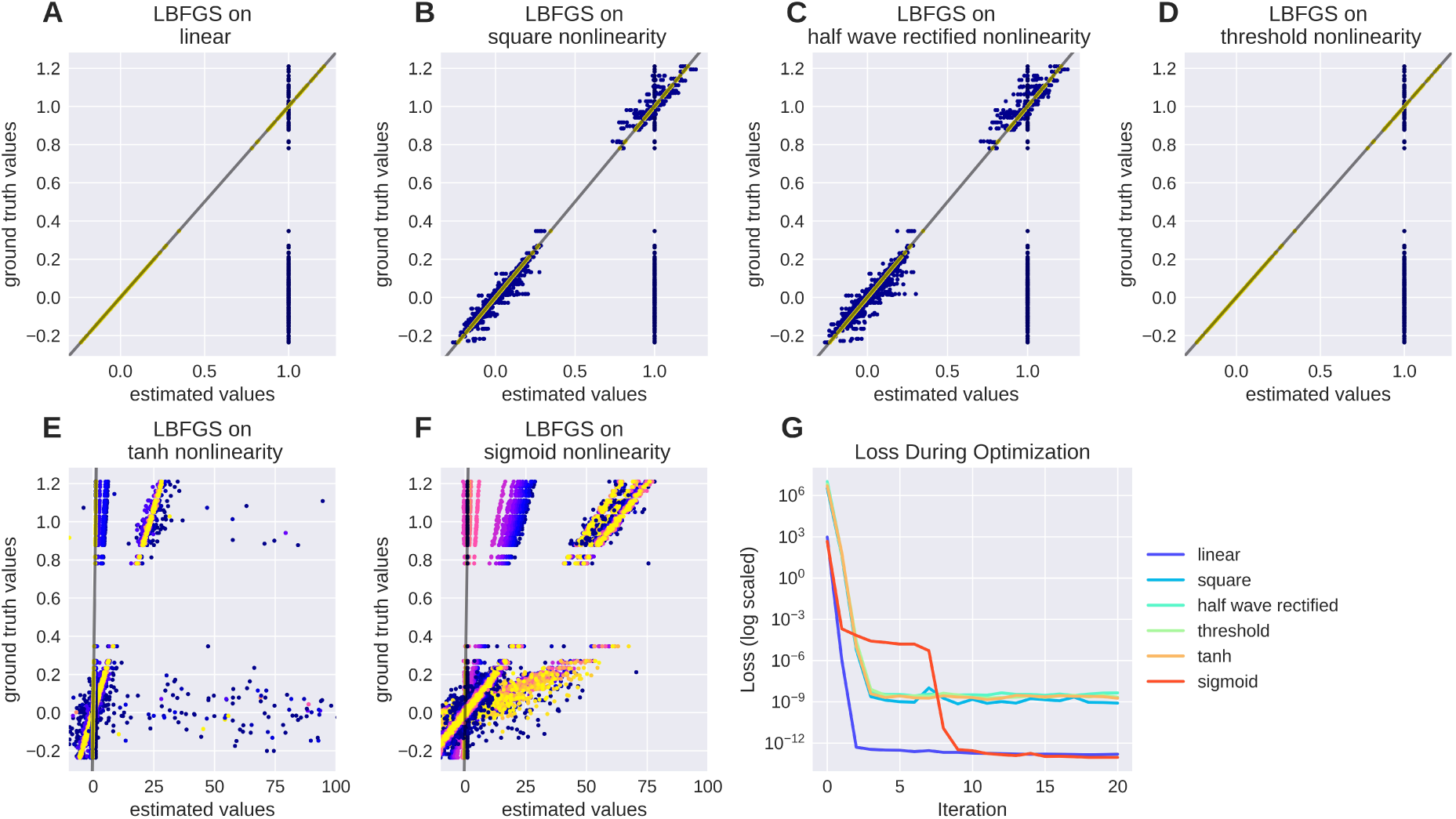
Recovering a receptive field under a range of non-linearities. LBFGS can find the solution after one step for linear and threshold non-linearity, very fast for squared output and half wave rectification and only after a larger number optimization steps, or a random re-initialization for tanh and sigmoid non-linearities.

#### 3.2.2 Usability Goals

Getting familiar with a new tool is always a time investment, especially for experimentalists. We chose to design Convis to be used in an interactive Python environment, even though this increases the learning curve compared to eg. a GUI centered application. The advantages of interactive Python sessions is that there is no additional configuration language to create models. Completely new models can be created and tested interactively, giving immediate feedback and existing models can be inspected and modified eg. by replacing non-linearities or by replacing a full numerical convolution filter with a recursive filter. One goal in the design of Convis is that during runtime all parameters and configuration options are discoverable - only a minimum of names should have to be memorized to use the software (eg. that models can be found under convis.models). All other names can be discovered by (1) tab completion in IPython [25], and other environments that inspect the attributes of Python objects, (2) printing a model or layer, revealing its layers and sub-layers and (3) accessing the doc strings of the underlying classes.

~~~
⋙ m = convis.models.LN()
⋙ m.<press tab>
⋙ print(m) # shows the structure of the model
LN(
(conv): Conv3d (1, 1, kernel_size=(1, 1, 1), stride=(1, 1, 1), bias=False)
)
⋙ help(m)
Help on LN in module convis.models object:
class LN(convis.base.Layer)
| A linear-nonlinear model with a convolution filter.
|
| Pads input automatically to produce output of the same size as the input.
|
| Parameters
| –––––
| kernel_dim: tuple(int,int,int) or tuple(int,int,int,int,int)
| Either the dimensions of a 3d kernel (time,x,y)
| or a 5d kernel (out_channels,in_channels,time,x,y).
| bias: bool
| Whether or not to include a scalar bias parameter in
| the linear filter
~~~

### 3.3 Implementation

#### 3.3.1 PyTorch as Backend

We chose to implement our toolbox in PyTorch to benefit from its just-in-time compilation and automatic differentiation mechanisms.

Just-in-time compilation allows for fast execution of complex operations, while still using the flexibility of the Python interpreter. PyTorch can trace the execution of a Python function and compile it efficiently using one of the compiler backends, combining native C or CUDA functions with on-the-fly created code to execute python for-loops as C for-loops.

Each computation that is performed is referenced in a computational graph attached to the output variables. These references can be used to compute values for each input variable that corresponds to the gradient of the output with respect to this input variable. These gradients can then be used in optimization algorithms to efficiently fit a model to data.

A discontinued Theano based version is still available at https://github.com/jahuth/convis_theano.

#### 3.3.2 Parameters and Modules

We build on the torch.nn module and provide a Layer class that adds a few convenience functions on top of the `torch.nn` Modules (see the documentation at https://jahuth.github.io/convis/). Similar to a `torch.nn.Module`, a Layer has to define two functions: an initializer `(__init__)`, which sets up all the variables and computational modules the Layer requires, and a function that does the actual computation `(forward)`. Parameters that are added as attributes to a Layer are automatically collected by PyTorch, such that optimization algorithms can find and optimize all available parameters of a model. Also the values of all variables of the Layer can be exported or imported. Since our models run not only once, but on a sequence of input, we need to distinguish between variables that signify a Parameter or a State. While a Parameter has a value that is either fix or being optimized, a State is time dependent and depends on the previous input. This can correspond to the last *n* time slices of input for convolutional kernels with length *n* in the time dimension or the *n* last input steps and *k* last output steps for recursively defined filters. When a model is first run on one input sequence and then on another, the States have to be reset when the input is changed. For the general behaviour of the model, only the Parameters are important and given the same input, the same model will produce the same States.

#### 3.3.3 Linear Filters

We provide two methods to apply a linear filter to every position of the three dimensional input: kernel convolutions and recursive filters. As mentioned before, they differ in how closely they can approximate the desired output. While recursive filters are bound to simplify responses (we implemented exponential decay filters in time and Gaussian filters in space), convolution filters can capture the contribution of each input pixel in a certain spatio-temporal window. Recursive temporal filters have few parameters, in the case of an exponential filter only the time constant, while convolution filters can have many hundreds. But when fitting these parameters, convolution filters can be estimated rapidly by efficient optimization algorithms, such as the Broyden-FletcherGoldfarb-Shanno method ([7], implemented in PyTorch as LBFGS), as can be seen in Figures 8 and 9. For the recursive filters we have implemented however, this specific method fails as can be seen in Figure 7. More basic gradient descent methods can still be used, but their convergence can be slow.

To be able to fit very slow processes with spatially variable receptive fields, we also implemented a hybrid filter (SoftConv) that multiplies a set of fixed recursive filters of increasing length with a set of spatial filters that can be optimized. These filters have a decreasing spatial resolutions with time and a smooth temporal profile, yet they can capture spatial details, are efficient to calculate (due to the temporal recursive filters) and to fit (due to the spatial convolution filters).

#### 3.3.4 Kernel Convolutions

In Convis, receptive fields can have an arbitrary shape. The receptive field of an RGC for example is comprised of an excitatory center and a suppressive surround. These two components also have different temporal dynamics making the overall receptive field space-time inseparable. While it is possible to construct it as the difference of two separable filters, Convis can use completely inseparable filters by instantiating a `convis.filters.Conv3d` filter and supplying a three-dimensional receptive field as a tensor. Non-separable filters can have any shape of receptive field and match eg. a pattern of motion (see Section 4.2). When fitting a model, a convolutional filter can be gradually adapted into a complex shape by gradient descent methods or even unsupervised learning and then analyzed post-hoc if it is possible to approximate it with more efficient filters.

#### 3.3.5 Recursive Filtering

Implementing an infinite impulse response filter for discrete time steps is not possible with a simple convolution kernel. So, in addition to convolutional filtering, we also reimplemented recursive temporal and spatial filters. While this restricts the shape of the filters, the computation is much more efficient, especially for long temporal filters. The parameters of the filter can be optimized using their gradient, which is an advantage over the implementation in VirtualRetina.

A temporal recursive filter relies on a finite number of past input and output values and multiplies them each with a vector of coefficients. The recursive filter we implemented is defined as:

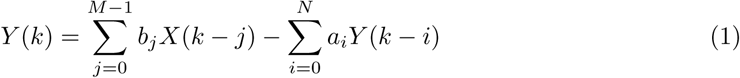

with *X* being the input and *Y* being the output. *M* is the number of the coefficients in vector *b*, *N* is the number of coefficients in vector *a*. For *N* = 0 the filter has finite impulse response. The computation has to be done in temporally correct order and the previous timesteps must be accessible. In contrast to convolution, which always only depends on a small, local area of the input and is independent of all previous computations, this is a challenge to parallelizing the computations of a recursive filter. In contrast to exponential filters, Gaussians are symmetric in both directions: If we interpret them as filters in time, they have a causal and an anti-causal component, in space they extend forward as well as backwards. A two dimensional Gaussian has an additional option for optimization: since a recursive definition in two directions simultaneously is very costly, splitting the computation into two stages that are computed independently and then combined improves the time complexity significantly. [11] uses for this the sum of a causal and an anti-causal filter while [34] use the product of a causal and anti-causal filter. The drawback of this optimization is that the Gaussians in x and y direction have to be independent, i.e. either a circular Gaussian or an ellipse whose major axis is aligned with either x or y. The recursive filters we implemented have exactly the same drawbacks as the ones implemented in VirtualRetina, except that it is possible to execute the computations on the GPU and gradients with respect to the filters parameters can be back-propagated.

## 4 Results

### 4.1 Simulating a Population of RGCs

To verify that our new implementation of the VirtualRetina model still replicates old results we made one-to-one comparisons using different stimuli and retina configurations.

Our implementation of the VirtualRetina model using Convis is available in the `convis.retina` submodule. To verify that the temporal responses are identical to the same model implemented in VirtualRetina, we used a full field chirp stimulus as used in [1] (see Figure 1) to characterize the temporal characteristics of retinal ganglion cells. The chirp features an OFF-ON-OFF pulse followed by oscillations increasing in frequency and then oscillations increasing in amplitude. The configuration was supplied as an xml file, as is typically used for VirtualRetina. Both simulations were configured with the same configuration file. Yet the Convis version created corresponding convolutional filters as opposed to recursive filers. As it can be seen in Figures 11,13, and 14, while the bipolar and ganglion cell stages replicate the results of VirtualRetina with high accuracy, the OPL layer has some numerical differences for very abrupt changes due to the low precision of the filters that are generated from the configuration. In Figure 12 we show a convolution kernel that is fitted to match the response of the OPL, showing that the response can be more faithfully reproduced by a single linear convolution filter. Overall, the output of the models is close enough that the difference is unobservable when spikes are generated and compared as either instantaneous rates or spike times. Figure 3 gives a quantification of the variance explained for each layer in isolation.

**Figure 10:**
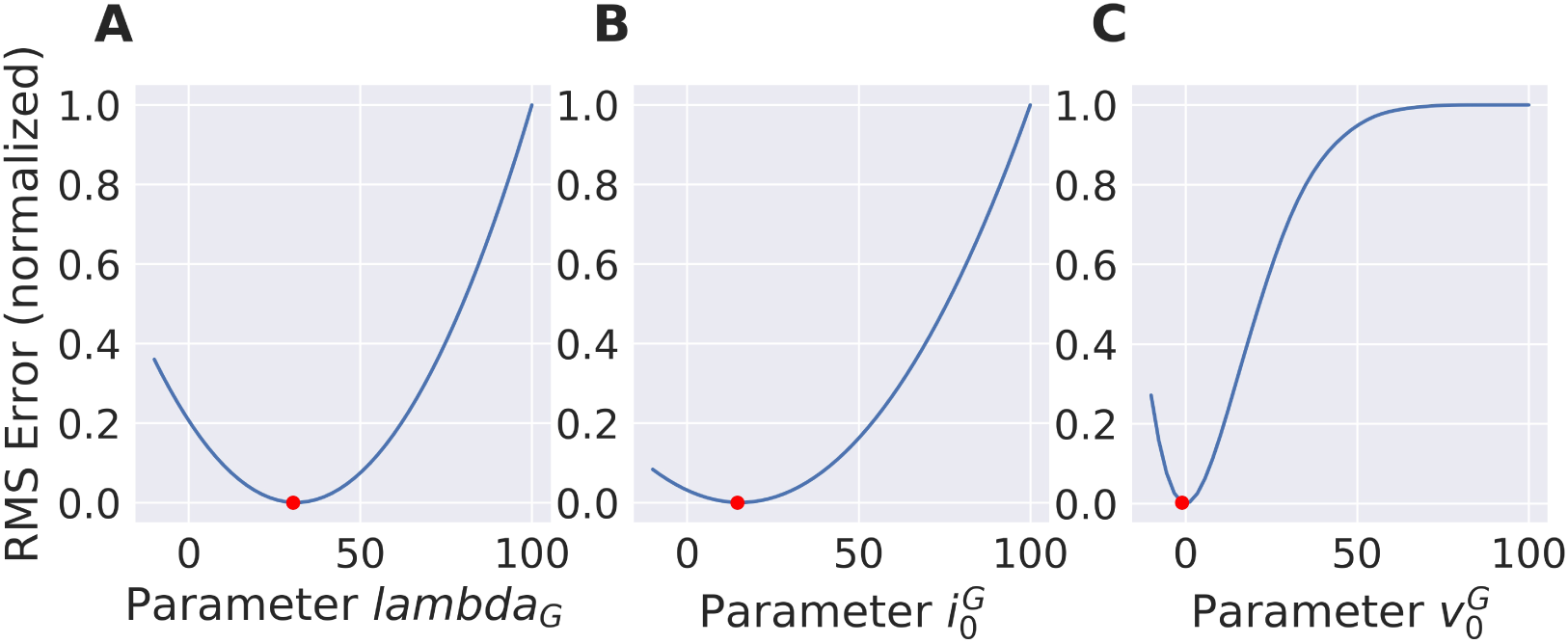
Nonlinear parameters do not show a squared error function. Still the shape is concave and the gradient points toward the minimum (red dot) which makes gradient based optimization very efficient.

**Figure 11:**
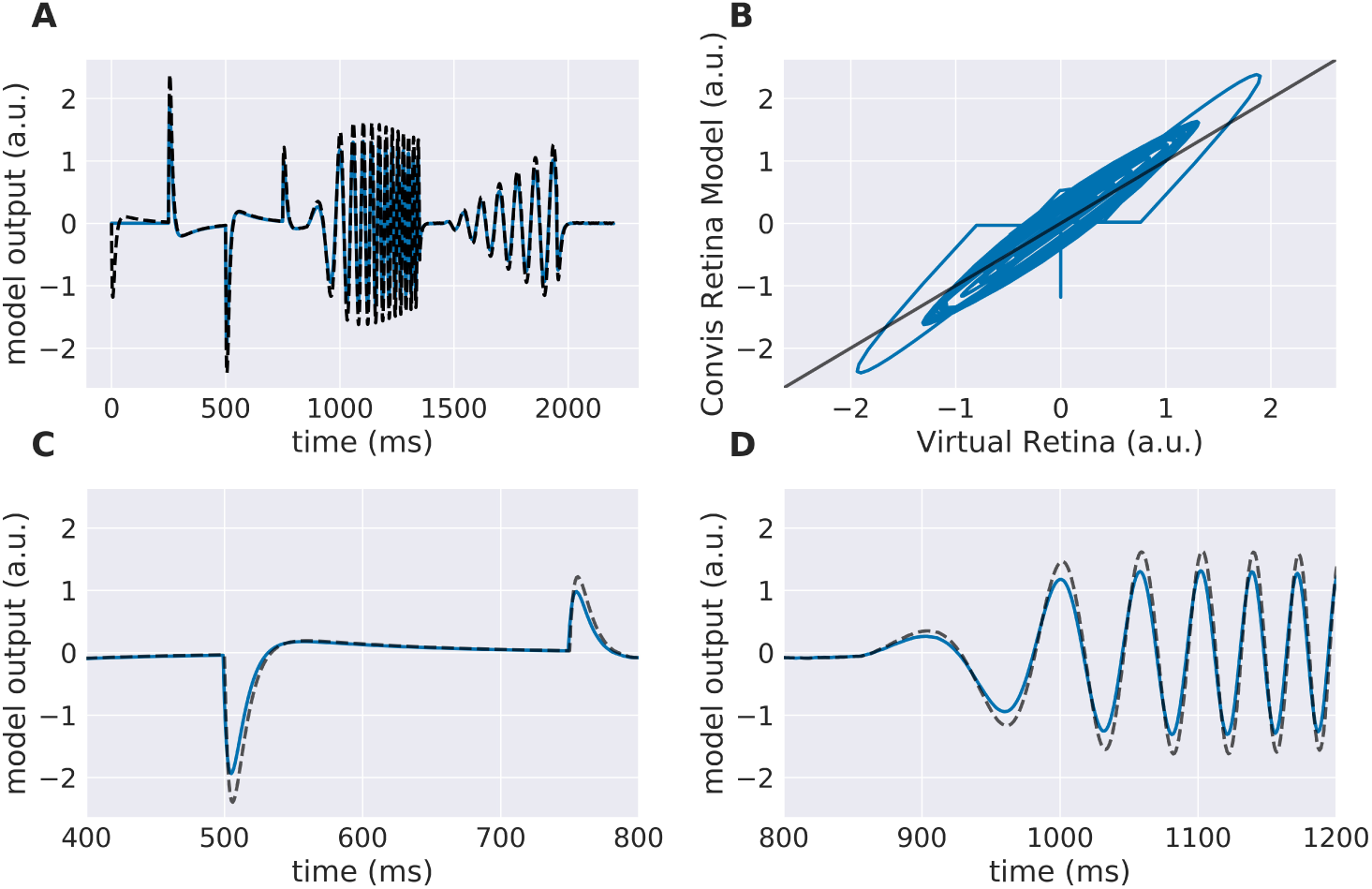
Comparison of Virtual Retina OPL stage (dashed line) and the OPL stage of the Convis Retina model (solid line): The stimulus is a “chirp” (see Figure 1). (A) shows the complete trial, (C) and (D) show details; (B) compares the trace of the original VirtualRetina to the new model. Due to numerical differences between the method of recursive filtering and convolutional filtering, the full-convolution OPL stage differs slightly from the original trace for very abrupt changes.

**Figure 12:**
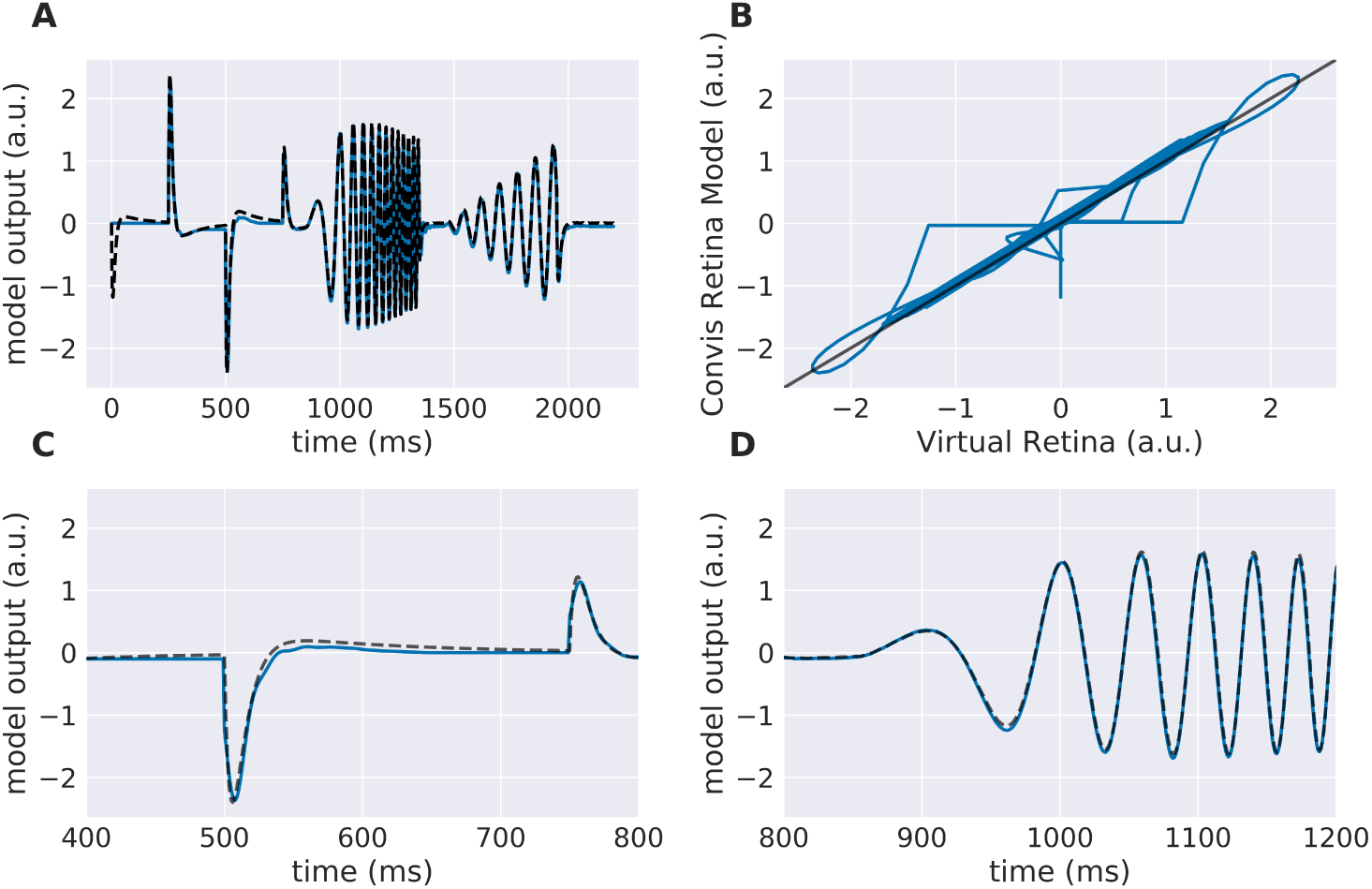
Comparison of Virtual Retina OPL stage (dashed line) and a linear model with a single convolutional filter (solid line): The OPL stage fitted to the desired response instead of using configuration values can reproduce the response of the VirtualRetina OPL faithfully. (A) shows the complete trial, (C) and (D) show details; (B) compares the trace of the original VirtualRetina to the new model.

**Figure 13:**
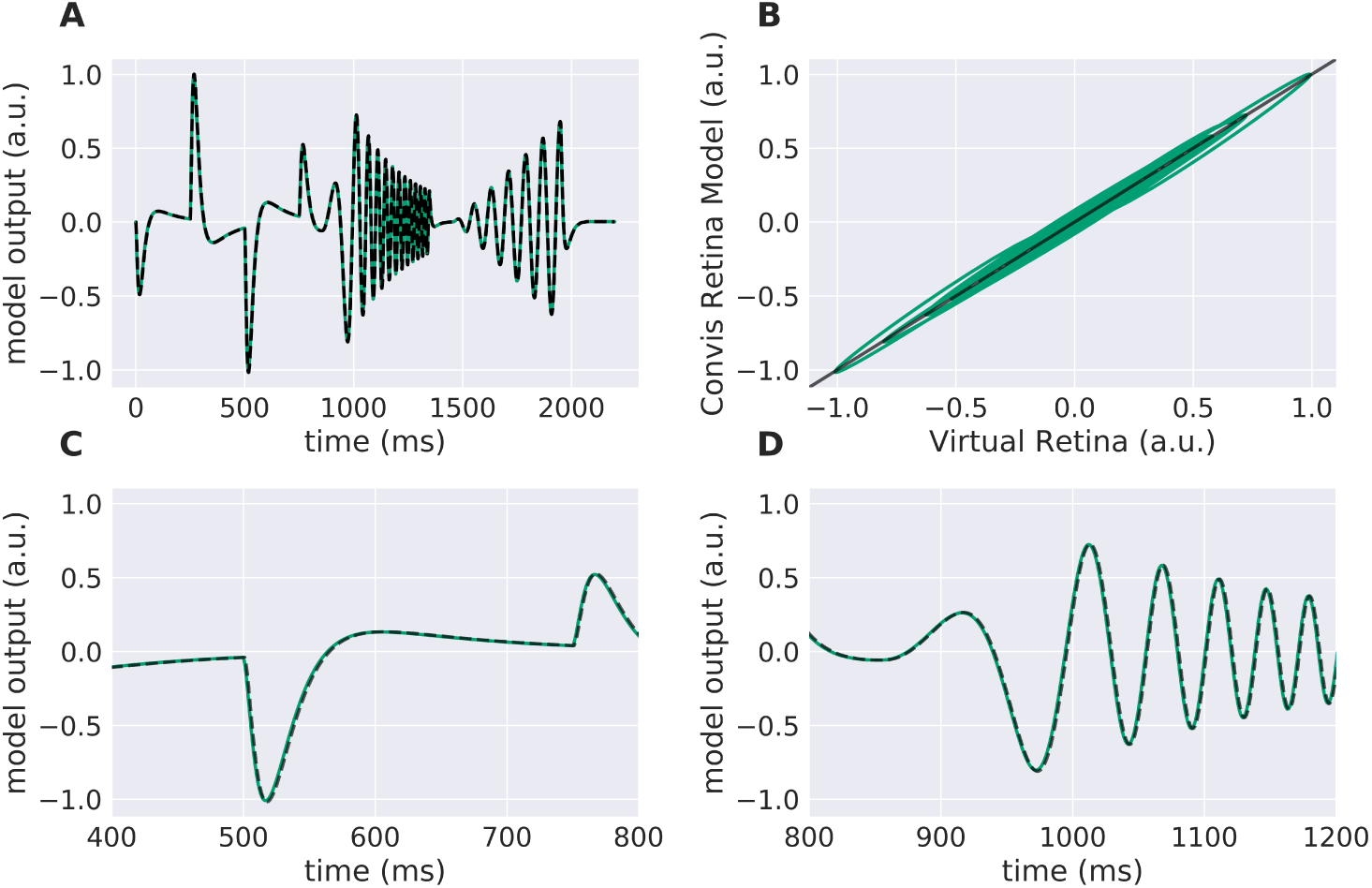
Comparison of Virtual Retina Bipolar stage (dashed line) and the Bipolar stage of the Convis Retina model (solid line). The stimulus is a “chirp” (see Figure 1). (A) shows the complete trial, (C) and (D) show details; (B) compares the trace of the original VirtualRetina to the new model.

**Figure 14:**
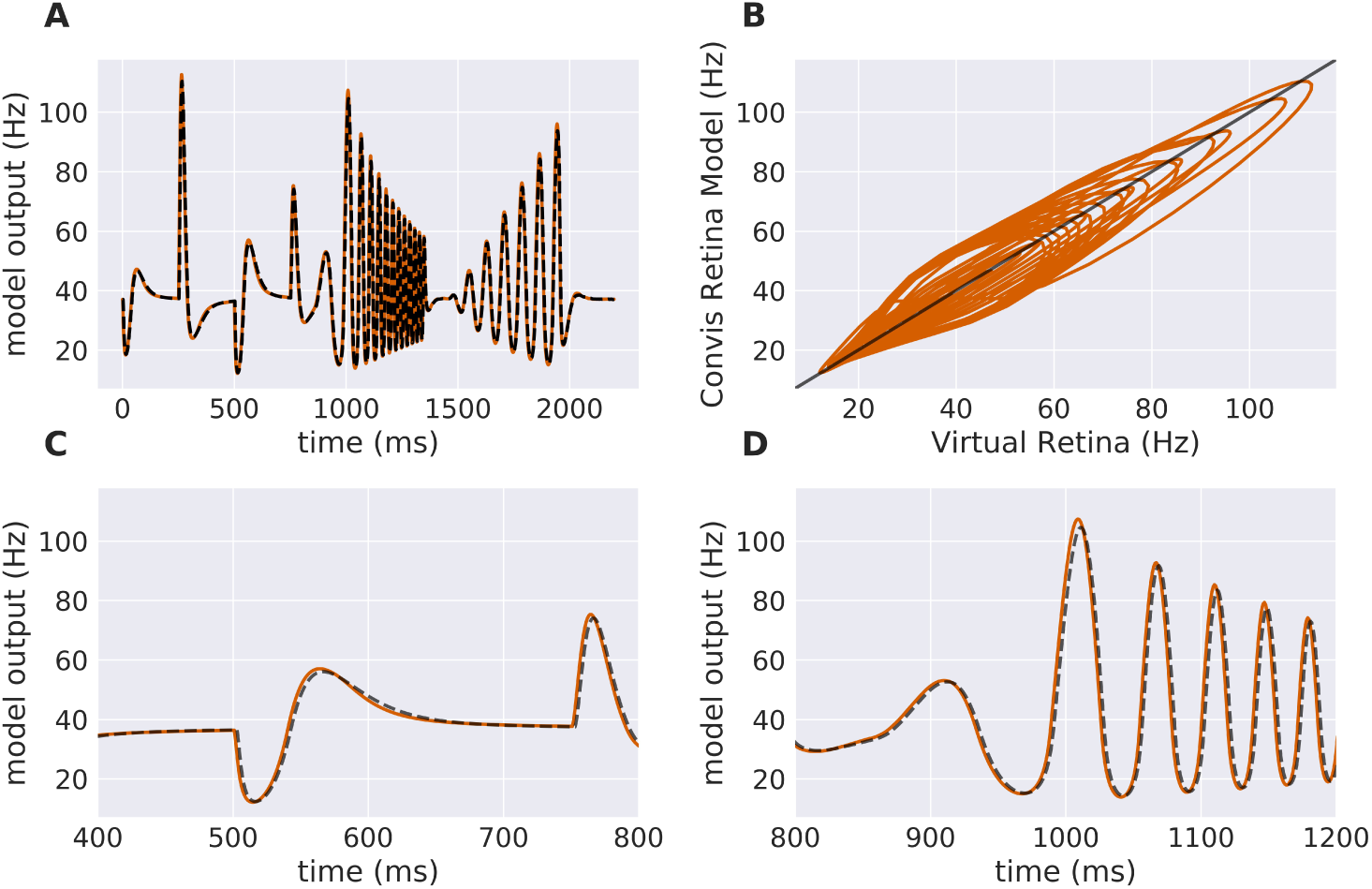
Comparison of Virtual Retina Ganglion Layer stage (dashed line) and the Ganglion Layer stage of the Convis Retina model (solid line). The stimulus is a “chirp” (see Figure 1). (A) shows the complete trial, (C) and (D) show details; (B) compares the trace of the original VirtualRetina to the new model.

### 4.2 Simulating Direction Selective Responses

The main advantage of full numerical 3d filters is that reactions to motion patterns can be captured in a straight forward manner. Movement sensitive receptive fields can be created with selectivity of certain speeds and directions, an example can be seen in Figure 4: A Gaussian that moves across the receptive field is most excited by a stimulus that travels in unison with it. To make the response ignore any other stimulus characteristics, a negative Gaussian closely follows the first, such that only an edge can excite the cell. To respond to both negative and positive edges, the absolute value of the response is taken. Figure 4 shows the resulting direction tuning curve.

These arbitrary filters can model a wide range of responses, including simple and complex V1 cells. Complex cells can be simulated either adding a rectification non-linearity and a second linear stage for spatial summation.

### 4.3 Estimating Parameters from Synthetic Data

#### 4.3.1 Fitting a spatial receptive field

Numerical filters have a large number of parameters, but since their behaviour is linear, they are easy to fit. To test that, we created a model with a convolutional 2d receptive field of a delicate structure: an image of the letters “con” and “vis” with some noise added. Figure 8 shows how well a simple gradient descent method can recover the shape of the filter from an linear-nonlinear model using a normal distributed noise stimulus. The SGD optimizer can find the parameters after some optimization steps while the `LBFGS` optimizer can jump right to the solution for simple nonlinearities (half wave rectification, thresholding The receptive field is even easier to recover for purely linear models or when the stimulus noise is more sparse. Figure 9 shows the `LBFGS` optimizer trying to estimate a linear filter that is subject to different non-linearities. While the optimization converges rapidly for simple non-linearities such as squaring or half-wave rectification, non-linearities with a very shallow gradient such as tanh or a sigmoid can pose a problem for the optimizer. Although a threshold non-linearity does not actually have an informative gradient at any point except the decision boundary, the PyTorch implementation of this operations backward function allows a fast convergence for this model (see Figure 9 D).

#### 4.3.2 Fitting a Recursive Exponential Filter

To show a simple example of optimization in the temporal domain we created a Poisson spike train which is convolved with an exponential filter of a certain time constant. Then we used the gradient with respect to the time constant of the recursively defined exponential filter to optimize this variable. As this is a fairly simple task, the only possible complication that can arise comes from choosing the learning rate parameter: convergence can be too slow or oscillate around the true value. This can be remedied by introducing a momentum term or adapting the learning rate over time. Figure 7 shows how the response of the model approaches the target. Again, naÏve gradient descent works, but it is not an efficient method as the direction of the gradient does not change, but the magnitude is proportional to the distance to the ground truth parameter and thus slows down the closer it is to the true value.

#### 4.3.3 Fitting Non-Linear Parameters

For deep-learning models, non-linearities are usually fixed to very simple functions without any free parameters. A half-wave rectification or a sigmoid function can be deemed parameter free, since their parameters concern the linear scaling of in - and output, which is already part of the linear filters of the model. In contrast, the VirtualRetina model has non-linear parameters that can not be emulated by scaling the linear filters. To describe the behaviour of experimental data, these parameters also have to be adapted to recreate the non-linear behaviour of the examined system. We examined the case that the examined system has the same general architecture as our model and only the value of the parameters differ. We evaluated the response of a ground truth model to a stimulus and then varied the parameters of a second model instance to examine the curve of the error function. The error function (see Figure 10) does not look like a square function, but still the function is concave and the gradient points toward the minimum, allowing gradient descent methods. It is possible to fit a polynomial to all points so far encountered, but the quality of the solution depends on the location in the parameter space and the noise in the fitting process. The most efficient way to find the true parameter is to get close enough that a 2nd degree polynomial is a good approximation to the local error function and then use Newton-like methods to find the optimum.

### 4.4 Using the Second Derivative

The gradients in Figure 8, C show that the gradients lie on a straight line. If the linear filter would have been subject to a stronger non-linearity, this would no longer be the case. The second derivative of the error with respect to a parameter will give rapid information about this. While in the simple case the 2nd derivative will be almost constant, it will vary more strongly for more complex non-linearities. Additional information about the derivatives of the error will decrease the number of samples needed to estimate the shape of the error function. As an example, in the simple case of a noise-less linear parameter, the error function is a 2nd degree polynomial. To estimate it, we would either need three samples of the error at different parameter values, or only two samples of the gradient, since we only have to fit a line rather than a parabola. If noise is present (as is the case in Figure 8), the advantage of estimating a lower-degree function can be even larger.

Another application of 2nd derivatives is to capture the interaction of parameters. Figure 17 shows the interaction between two linearly interdependent parameters in the Bipolar stage of the retina model, given a random checker-board flicker stimulus and a ground-truth model with default parameters. We show the gradient with respect to each parameter and a resulting flow field over a visualization of the error. A long corridor of almost identical solutions exists on the line g_leak_ = λ_bip_. From this plot we can already assume that gradient descent will be very inefficient, but using a Hessian based descent method is more effective.

**Figure 15:**
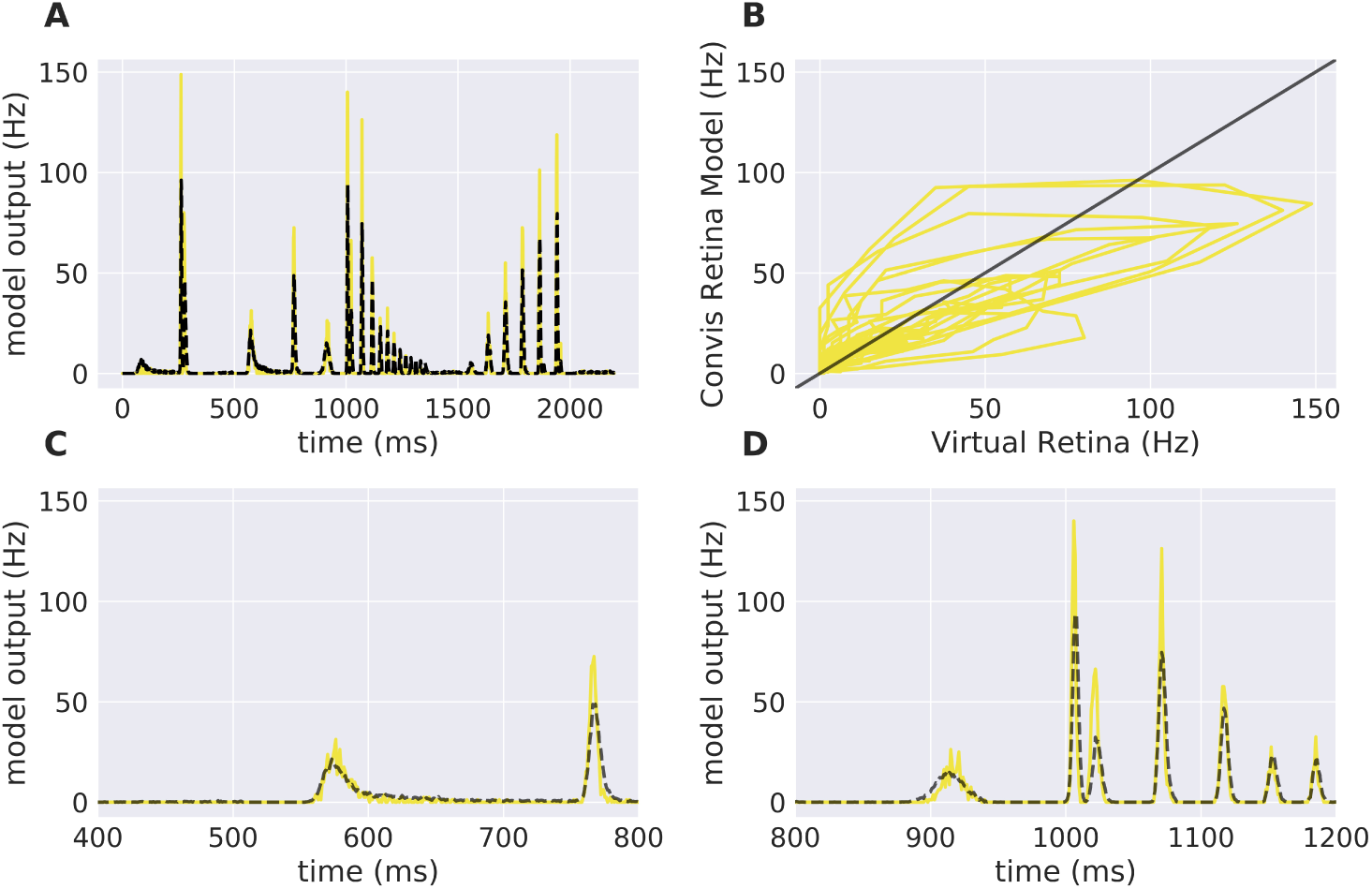
Comparison of Virtual Retina Ganglion On Spiking Layer stage (dashed line) and the GanglionSpiking stage of the Convis Retina model (solid line). The stimulus is a “chirp” (see Figure 1). (A) shows the complete trial, (C) and (D) show details; (B) compares the trace of the original VirtualRetina to the new model.

**Figure 16:**
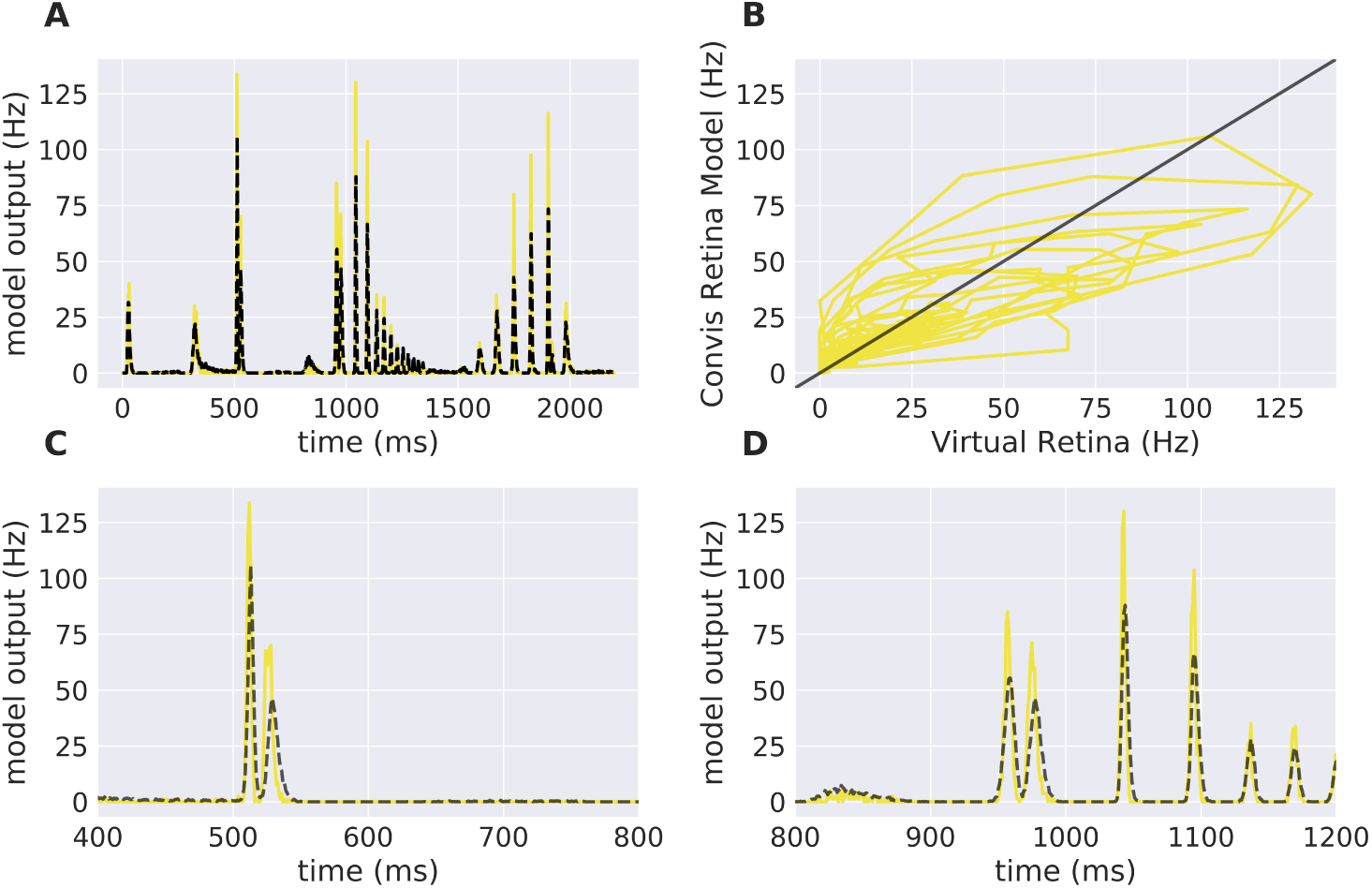
Comparison of Virtual Retina Ganglion Off Spiking Layer stage (dashed line) and the GanglionSpiking stage of the Convis Retina model (solid line). The stimulus is a “chirp” (see Figure 1). (A) shows the complete trial, (C) and (D) show details; (B) compares the trace of the original VirtualRetina to the new model.

**Figure 17:**
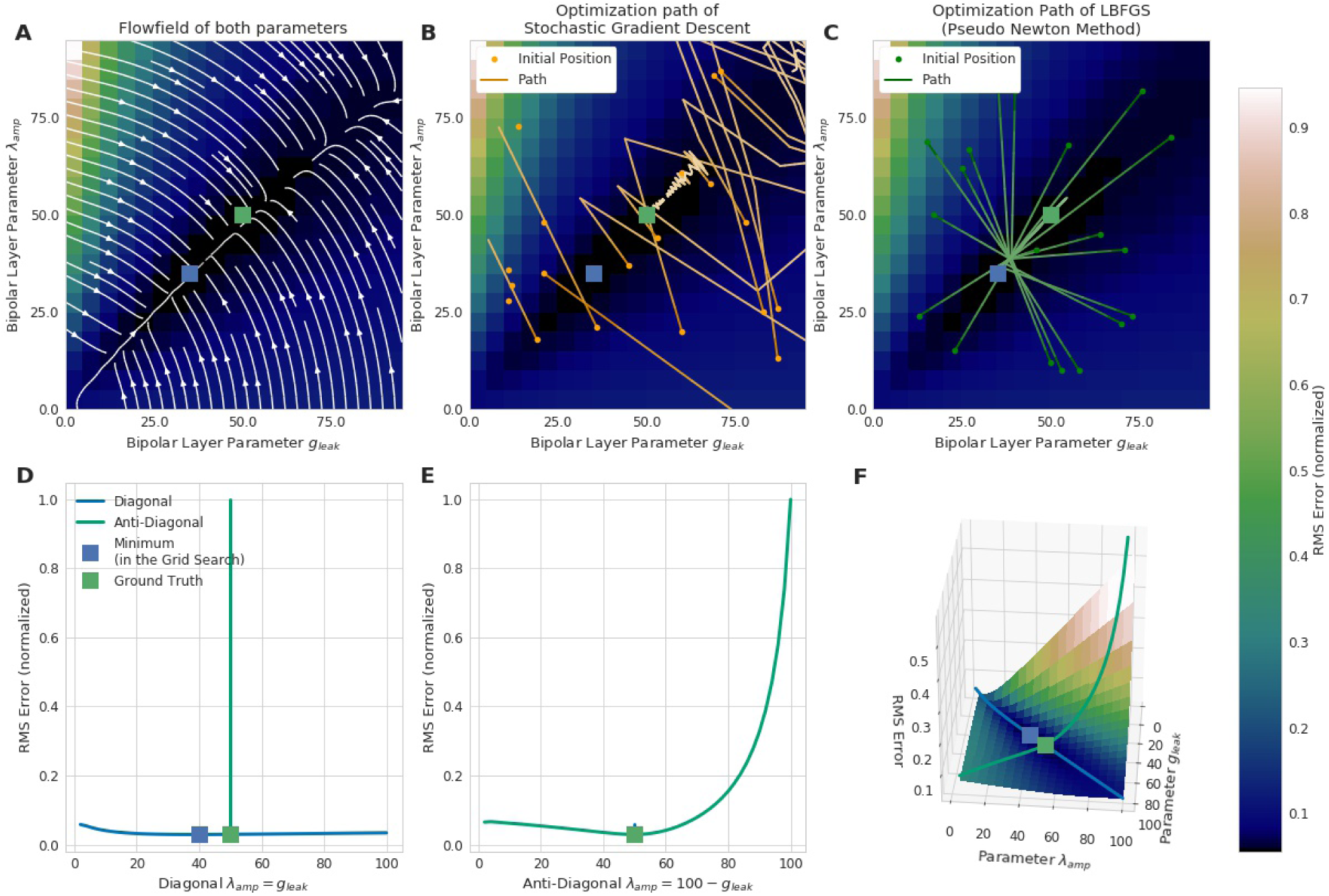
The interaction of two parameters in the Convis Retina model poses a problem for naive optimization methods. We visualized a grid search for the true parameters of a ground-truth model in terms of the gradients with respect to each of the parameters and the error function with an overlay of the flow field resulting from the two gradient functions. The blue square shows the global minimum of the grid search (which does not correspond to the ground truth values of the two parameters since the parameters lie between the grid as can be expected when parameters are unknown). (A) shows the combination of both gradients. Most gradients do not point directly at the true minimum or at the ground truth, but rather at the a “valley” along the diagonal. (B) and (C) show the path of two optimizer algorithms when searching for the parameters. The stochastic gradient descent method has trouble finding the minimum since the direction of the gradients carry only little information. The optimizer “bounces” between the walls and started to escape to increasingly large values (trials were stopped once the parameters left a certain region). The pseudo Newton method of the LBFGS shows a quick convergence towards one point in the valley and a subsequent approach to the true values. So it can be noted that although the error plane is convex, grid search found a minimum, which was not very close to the true minimum in the parameter space, gradient descent had great difficulties finding even the valley. Gradient descent methods need very sophisticated learning rate schedulers to adapt to the large range of gradients. LBFGS converged to the same point independent of initial condition, which is due to the self guided exploration the methods uses to approximate the error surface. But to get closer to the true parameters, more iterations were needed. (D) and (E) show the profile of the diagonal and anti-diagonal (as shown in (F)). The error along the diagonal is much shallower than the gradients on both sides of the valley. (F) shows the error in a surface plot with the diagonal and anti-diagonal highlighted.

### 4.5 Optimizing the Input to a Model

We set up a small simulation in which we simulated two retina models (A and B) and took the difference of their outputs. The configuration between the models differed slightly. Then we created a moving grating stimulus, dependent on a few parameters, such as direction and speed. We added this stimulus as an input to the model, building a graph of operations. When we now vary the speed of the gratings, the difference between the two models will change, for one, there is only a certain range where the moving gratings are actually visible for the model due to bandpass filtering (and the limitations of resolution). Figure 5 shows the optimal speed to discern the two models. The function is not a simple relation, and even shows more than one local maximum, possibly due to harmonics. We can compute the gradient of the difference with respect to the speed parameter of the gratings, which fits well with the actual error observed. If this were an actual experiment and we were trying to decide whether model A or B is closer to the actual mechanism we were observing, we could now choose the stimulus that will give us the greatest amount of information.

An application of this method could be the creation of tailored stimuli to distinguish a set of a-priori known classes of cells. Stereotypical responses shown eg. in [1] can be used to fit a prototype model per class. Next a parametrized function that generates a visual stimulus can be optimized for a pair of models each such that when that specific stimulus is presented, the output of the two models is maximally different. A sequence of these optimal stimuli can then be used to rapidly classify cells into known classes, even if their responses eg. to a chirp stimulus is very similar, which would require a large number of trials to reliably classify a cell using current methods of eg. estimating and matching the temporal profiles of the receptive fields. A different application concerns the problem of model fitting and validation itself. Depending on the exploration of the parameter space and the complexity of the model, multiple parameter combinations can achieve a similar fit to the data. These sets of solutions might behave similar on the observed stimulus, but different on stimuli that were not observed. Finding these stimuli can be posed as an optimization problem on a parametrized input stimulus.

It is possible to do any of these computations online, during an experiment. Depending on the complexity of the model, the fitting process will introduce a lag (see Section 4.6.1 for an estimate of running speed), however given the possibility of parallelization and the decreasing costs of GPUs, repeated model fitting and stimulus optimization during an experiment is achievable.

### 4.6 Comparison to Other Simulation Software

As we have shown in section 4.1, the response of Convis is identical to the VirtualRetina if an identical configuration is used. The linear filtering and gain control properties are identical within numerical errors the magnitude of which is configurable. In addition to those response characteristics, Convis can also simulate non-circular, non-Gaussian and non-continuous receptive field shapes. Similarly to COREM, Convis can be configured more flexibly, combining basic filters.

Table 1 shows a comparison between different retina simulation software.

**Table 1:** Comparing different retina simulation software: VirtualRetina [35], Topographica [4], Virtual Retina++ (continued development of VR in the ENAS / PRANAS package [9]), Lorach et al. [17], and RetinaStudio [19]. ≈VR signifies that the gain control was implemented similarly to VirtualRetina: Contrast gain control through shunting inhibition, using a local estimate of spatio-temporal contrast [36].

| Name / Application | Luminance Gain Control | Contrast Gain Control | Language of Source Code, Configuration | Open Source | Filters | continuous in/output | Optimization/ Plasticity |
| --- | --- | --- | --- | --- | --- | --- | --- |
| VirtualRetina [35]<br>RGC responses | Yes | Local Shunting | C++, xml | Yes (CeCILL-C) | recursive | No | No |
| Topographica [4]<br>PVC / Neural Maps | Yes | Naka-Rushton [24] | C, Python | Yes (BSD 3) | convolution |  | Yes |
| ENAS [9]<br>Model Verification | Yes | ≈VR | C++, xml/GUI | No | recursive | No | No |
| Lorach et al.<br>Neuroprosthetics | Yes | No | hardware DVS, Matlab | No | convolution | No | No |
| COREM [20]<br>RGC responses | Yes | ≈VR | C++, scripts | Yes (CeCILL-C) | recursive | Yes | No |
| RetinaStudio [19]<br>Neuroprosthetics |  |  | C#/Flowlang | Flowlang |  | No | No |
| isetbio [6]<br>Perceptual thresholds optical aberrations photoreceptor sampling |  | - | Matlab / | Yes (MIT) | hexagonal grid | No | No |
| pulse2percept [5]<br>Perceptual thresholds of RGCs for implant assessment |  | - | Python / Scipy | Yes (BSD 3-clause) | square or gaussian RFs, radial current spread | No | No |
| iModel Ret_Mesh [2]<br>RGC responses | Yes | No | C/OpenGL / Java | code available from imodel.org website | hexagonal grid RFs | No | No |
| deepretina [23]<br>RGC responses | HP filters possible | No | Python / Theano | available on github (no license specified) | convolution | No | Yes |
| Convis<br>RGC responses | Yes | ≈VR | Python / PyTorch | Yes (GPL-3) | recursive or convolution | Yes | Yes |

#### 4.6.1 Calculation Speed

For the application in closed-loop experiments, running time can be crucial. The additional features of Convis compared to VirtualRetina come at a cost of calculation speed. The computational graph adds a small overhead to each operation (in the order of *10μS* according to the PyTorch developers). Convolutions are also remarkably slower than recursive filtering for large filters. Recursive circular Gaussian spatial filtering is separable into *x* and *y* components, which can be efficiently computed while convolutions have to process *x_image_* × *y_image_* × *x_filter_* × *y_filter_* single multiplications. Including a time dimension into the filters which are not separable further increases the complexity drastically. Even when these calculations are done on a GPU, there is some overhead in starting the computations.

To assess how useful Convis can be for specific applications, we compared VirtualRetina and a set of Convis models over a range on stimulus sizes (see Figure 6). The comparison was done on a Dell PC with a 6-core Intel(R) Xeon(R) CPU E5 v3 running at 2.40GHz, 32gb of RAM and a NVIDIA Quadro K620 GPU. For all models small stimuli could be processed faster than real-time and larger stimuli increased the running time polynomially. The Convis Retina model is for small stimulus sizes the slowest model and also has the smallest stimulus size that still allows real-time processing (20x20 pixel on our machine). The original VirtualRetina program outperformed the Convis model for all stimulus sizes and could still process 40x40 pixel images in close to real-time. However, this is only the case if the only relevant output are the produced spikes. When the input current to the ganglion cells is required for further analysis, the data has also be written to a hard disk which brings the overall running time close to the Convis Retina model.

For a use case where just spikes are required, receptive fields can be assumed circular and the model need not be optimized, VirtualRetina has a strong speed advantage over the Convis Retina model. If however the model has to be fitted to data, the speed increase through gradient based optimizations compared to gradient-free optimizations is reversed. If non-circular receptive fields are required, VirtualRetina is not applicable at all.

The linear-nonlinear convolution models - LN, LNLN and a convolution model similar to [23] - show a very similar increase in computation time: They are faster than all other models for very small input stimuli and slower than all other models for large stimuli sizes due to the computational complexity of 3d convolution.

#### 4.6.2 Features of VirtualRetina not Implemented

We did not implement code to parse radially varying parameters. While the general idea of retinotopic varying parameters can be useful, we found that we were interested in rather local retinal populations (< 2 deg) which will be fairly uniform. Radially varying blur can still be used when filtering with recursive spatial filters, but the configuration values for radially varying blurs in VirtualRetina xml files will be ignored by default. Also, we did not implement cell sampling schemes. One pixel is one cell for us, however selecting a subset of cells from the output (before compiling the model) is possible and it can reduce the computations actually performed by the model due to the optimization mechanisms.

## 5 Discussion

Using the Convis toolbox we can create models for responses to visual stimuli, eg. retinal ganglion cells or V1 cells.

Ganglion cell responses can be simulated comparably to VirtualRetina on an equally large scale (Section 4.1) while being able to switch between Gaussian receptive fields and arbitrary receptive field shapes. If users of VirtualRetina want to use receptive fields with arbitrary shapes or modify the model on the fly, Convis offers that possibility.

For a more general approach, Convis includes more general linear-nonlinear models. To fit them to motion sensitive cells, Convis can use non-separable convolution filters to implement filters in a very straightforward manner (Section 4.2) which can be optimized by a range of established, gradient-guided methods. Large, linear convolution filters also have the advantage that their parameters, although numerous, are easy to fit (section 4.3). Still, non-linear parameters can be optimized with the same methods (4.3.3).

The open architecture of PyTorch and thus Convis makes it easy to build custom models and build on traditional LN models, extending them with machine learning tools such as recurrent or deep neural networks, as was done by [3] using a Theano.

The stimulus plays an important role when comparing a model and a physical system. To aid experimenters in choosing a good stimulus, we demonstrated that also the parameters of a stimulus can be optimized with gradient descent to maximize the difference between two very similar models and thus the information that new experiments will provide for the model fitting process (Section 4.5).

## Conflict of Interest Statement

The authors declare that the research was conducted in the absence of any commercial or financial relationships that could be construed as a potential conflict of interest.

## Funding

This research was supported by ANR - Essilor SilverSight Chair ANR-14-CHIN-0001.

## Acknowledgments

Thanks to Adrien Wohrer for helpful insights into the workings of VirtualRetina and thanks to Richard Carrillo for insightful comments.

